# Primary cilia deficiency in neural crest cells causes Anterior Segment Dysgenesis

**DOI:** 10.1101/752105

**Authors:** Céline Portal, Peter Lwigale, Panteleimon Rompolas, Carlo Iomini

## Abstract

During eye embryogenesis, neural crest cells (NCC) of the periocular mesenchyme (POM) migrate to the anterior segment (AS) of the eye and then differentiate into the corneal stroma and endothelium, ciliary body, iris stroma, and the trabecular meshwork. Defective development of these structures leads to anterior segment dysgenesis (ASD) that in 50% of the cases leads to glaucoma, a leading cause of blindness. Here, we show that the primary cilium is indispensable for normal AS development and that its ablation in NCC induces ASD phenotypes including; small and thin cornea, impaired stromal keratocyte organization, abnormal iridocorneal angle with reduced anterior chamber and corneal neovascularization. These defects are similar to those described in patients with AS conditions such as Axenfeld-Rieger syndrome and Peter’s anomaly. Mechanistically, disruption of the primary cilium in the NCC resulted in reduced hedgehog (Hh) signaling in the POM, canonically activated by the Indian Hedgehog ligand expressed by endothelial cells of the choroid. This caused decreased cell proliferation in a subpopulation of POM cells surrounding the retinal pigmented epithelium. Moreover, primary cilium ablation in NCC also led to a decreased expression of *Foxc1* and *Pitx2*, two transcription factors identified as major ASD causative genes. These findings suggest that primary cilia are indispensable for NCC to form normal AS structures via Hh signaling. Defects in primary cilia could, therefore, contribute to the pathogenesis of ASD, and to their complications such as congenital glaucoma.

## INTRODUCTION

Anterior segment dysgenesis (ASD) is a term referring to a spectrum of congenital disorders of structures of the anterior segment (AS) of the eye. Abnormalities typically associated with ASD include; corneal opacity, cataract, posterior embryotoxon, iris hypoplasia, corectopia or polycoria, adhesions between iris and cornea or lens and cornea [1]. Approximately 50% of ASD patients develop glaucoma that can lead to visual impairment and blindness [1–4]. Although mutations in genes encoding transcription factors, transporters, and glycosylating proteins have been described in patients with ASD, several ASD cases still await genetic elucidation [1–4]. Moreover, the cellular and molecular mechanisms underlying the pathogenesis of different ASD conditions remain unknown. Most AS structures are derived from the neural crest cells (NCC), including the corneal stroma and endothelium, ciliary body muscle and body, iris stroma, and the trabecular meshwork [5]. In mice, migrating NCC of the periocular mesenchyme (POM) begin to invade the AS of the eye between E11.5 and E12.5, and differentiate into corneal endothelial cells and keratocytes[6, 7]. By E16.5, the presumptive iris is visible and detaches from the cornea, while the drainage structures (trabecular meshwork and Schlemm’s canal) continue to develop postnatally [8]. NCC also give rise to the sclera, pericytes of the choroid and hyaloid vasculature, orbital cartilage and bone, and oculomotor tendons [9]. The neural crest is a transient embryonic structure in vertebrates that delaminates from the border between the neural plate and the non-neural ectoderm; NCC migrate throughout the embryo to multiple locations and differentiate into a wide variety of cell types and tissues [10, 11]. Recent studies have shown that primary cilia play essential roles in morphogenetic processes involving neural crest-derived cells such as craniofacial development [12, 13]. It has been proposed that primary cilia seem to mediate tissue-tissue interactions requiring reciprocal signaling rather than purely NCC specification [14, 15].

Primary cilia are microtubule-based cellular organelles that emanate from the basal body and extend from the plasma membrane. A bidirectional movement of protein particles along the axoneme, called intraflagellar transport (IFT), ensures the appropriate assembly and maintenance of cilia [16–20]. Primary cilia play a pivotal role in the development and tissue homeostasis by regulating the Hedgehog (Hh), Wnt, and Notch signaling pathways among others [21, 22]. Hh is the pathway the most strongly associated with the primary cilium [23, 24]. The binding of one of the three mammalian Hh ligands (Sonic hedgehog (SHH), Indian hedgehog (IHH) or Desert hedgehog (DHH)) on the receptor Patched (PTCH) induces the exit from the cilium of PTCH and an accumulation of the Hh transducer Smoothened (SMO) in the cilium. Subsequently, SMO activates the Gli transcription factors which translocate in the nucleus and activate the expression of Hh target genes [25]. Target genes of the Hh pathway are involved in cell proliferation, maintenance of stemness, cell-fate determination, cell survival and epithelial to mesenchymal transition [26].

Dysfunctions of cilia lead to a wide range of human diseases called ciliopathies, that affect most human organ systems [24]. Conditions of the AS, including ASD, were described associated with ciliopathies. Patients affected by Meckel syndrome, a severe ciliopathy, present AS abnormalities including microphthalmos/anophthalmos, aniridia, cryptophthalmos, sclerocornea, abnormal corneal thickness, corneal neovascularization, abnormal iridocorneal angle, tunica vasculosa lentis, and persistence of nuclei in lens fibers [27]. Cases of microphthalmia were recently reported in patients affected by oral-facial-digital syndrome [28, 29]. Corneal opacity has been detected in a patient affected by Joubert syndrome [30], glaucoma and cataract are common conditions associated with Bardet-Biedl and Lowe syndromes [31, 32], and cataract and keratoconus with Leber’s congenital amaurosis [31]. Moreover, one of the features of the Biedmond syndrome type 2, resembling to the Bardet-Biedl syndrome, is the presence of iris colloboma (OMIM #210350).

Several studies focusing on craniofacial development have demonstrated that ablation of the primary cilium via inactivation of the IFT in NCC leads to severe craniofacial defects [14, 33–37]. Interestingly, systemic mutations in ciliogenic genes causing an increase of the Hh activity and a deletion of *Gli3*, which acts predominantly as a repressor of the Hh target genes, lead to similar abnormal ocular development [38–42]. On the other hand, the loss or downregulation of the Hh activity leads to severe craniofacial and ocular defects including anophthalmia, cyclopia, microphthalmia, and coloboma [9]. This leads us to hypothesize that the primary cilium plays a pivotal role in the development of the AS and that dysfunction of the primary cilium in NCC could lead to conditions similar to those associated with ASD.

In this study, we investigated the role of the primary cilium in the development of the NCC derived ocular structures and its possible role in ASD. We showed that NCC of the POM are ciliated and that primary cilia persist in keratocytes of adult mice. We demonstrated that ablation of the primary cilium in NCC induces an ASD phenotype with impaired corneal stroma organization and dimension, abnormal iridocorneal angle and corneal neovascularization.

We also showed a reduction of the Hh signaling pathway specifically in a subset of cells in the POM surrounding the retinal pigmented epithelial cells (RPE). Furthermore, we identified the endothelial cells of the choroid as the cells expressing *Ihh*, which is the Hh ligand maintaining the signaling activity in the POM surrounding the RPE layer in normal eye development.

## RESULTS

### Primary cilium ablation in NCC leads to ASD phenotype

To determine spatiotemporal distribution of primary cilia in NCC-derived tissue of the POM we visualized cilia by immunofluorescence (IF) using an anti-Arl13b Ab, a widely accepted ciliary marker [43] in genetically labeled NCC. NCC were traced by using the *Rosa26^mT/mG^*(mT/mG) reporter mouse line [44] crossed to the *Wnt1-Cre* mouse [45] in which the Cre recombinase is under the control of the *Wnt1* promoter, a gene highly expressed in early stages of NCC specification. In the *Wnt1-Cre;mT/mG* transgenic line, the Cre-dependent excision of a cassette expressing the red-fluorescent membrane-targeted tdTomato (mT) allows the expression of a membrane-targeted green fluorescent protein (mG) in bona fide NCC-derived tissues (**Figure 1A**). We observed that at E14.5, all NCC-derived tissues of the POM and the presumptive corneal stroma were ciliated (**Figure 1A**). Our previous studies reported that while primary cilia are present in developing corneal endothelium (also a NCC-derived tissue), they disassemble in adult corneal endothelium at steady state [46]. To assess the presence/absence of primary cilia in adult corneas we utilized a transgenic mouse line expressing the ciliary membrane protein somatostatin receptor 3 fused to GFP under the ubiquitous promoter for actin (Sstr3::GFP) [47]. Intravital microscopy revealed that cilia were present in all keratocytes of the corneal stroma of 3-month-old mice (**Figure 1B**). To gain ultrastructural insights we analyzed corneal stroma and POM in developing and adult eyes. TEM showed that in developing eyes, cilia emanated from the cellular surface into the extracellular matrix whereas, cilia of newborn keratocytes appeared to be intracellular or largely invaginated in a long ciliary pocket with their axis parallel to the cell plane (**Figure 1D-E**). Interestingly, the tip of cilia in developing corneas and POM were observed to interact with cellular protrusions of the neighboring cells (**Figure 1C-D**). Moreover, the plasma membrane of the cellular protrusion at the contact point with the ciliary tip appeared to be highly electron-dense, suggesting the presence of protein components or modified lipids in this region (**Figure 1D-E**). To investigate the involvement of primary cilia in AS development we set out to ablate *Ift88*, a subunit of the IFT machinery required for cilia assembly and maintenance [48], in NCC. To do so, we generated the *Wnt1-Cre;Ift88^fx/fx^* mouse (cKO) which was indistinguishable from the null hemizygous *Wnt1-Cre;Ift88^fx/-^*. In this mouse the *Ift88* gene is excised in all migrating mesenchymal cells expressing *Wnt1* leading to complete ablation of the primary cilium (**Supplement Figure 1A**) [45, 49]. To monitor ablation of cilia in the NCC of the POM we generated the *Wnt1-Cre;IFT88^fx/fx^;mT/mG* mouse and we labeled cilia with an anti-Arl13b Ab [43]. In control mice, virtually all the NCC of the POM appeared ciliated during development (**Fig. 1A**). In contrast, in the *Wnt1-Cre;IFT88^fx/fx^;mT/mG* mutant mice cilia were absent in most of the POM cells expressing Cre (**Supplement Figure 1B**). We confirmed ablation of cilia in keratocyte precursors by TEM. In keratocyte precursors, the basal body apparatus was observed to reach the apical plasma membrane in both control and mutant corneas, however, primary cilia were only observed emanating from basal bodies of control keratocytes (**Supplement Figure 1C**). *Wnt1-Cre;Ift88^fx/fx^*mutant mice died at birth and E18.5 embryos displayed strong craniofacial defects including increased frontal width, wider frontonasal prominence and increase of the distance between the nasal pits, consistent with previous studies (**Figure 2A**) [36, 37]. In addition, we detected abnormalities in the anatomy of the eye. At E14.5, a developmental stage preceding the closure of the eyelids, the axis of the eyeballs was misaligned facing downward in the cKO whereas, in control it remained perpendicular to the sagittal plane of the head (**Figure 2A**). Furthermore, the outline of the presumptive iris appeared irregular in the mutant while in the control it described a nearly perfect circle (**Figure 2A**). Later in development (E18.5), the irregularities of the iris exacerbated and the developing cornea appeared smaller than that of the control, strongly arguing toward severe defects of the AS including a reduced anterior chamber (**Figure 2**). Thus, these macroscopic anatomical abnormalities suggest a morphogenetic condition of the AS in the eye of the mutant.

**Figure 1:**
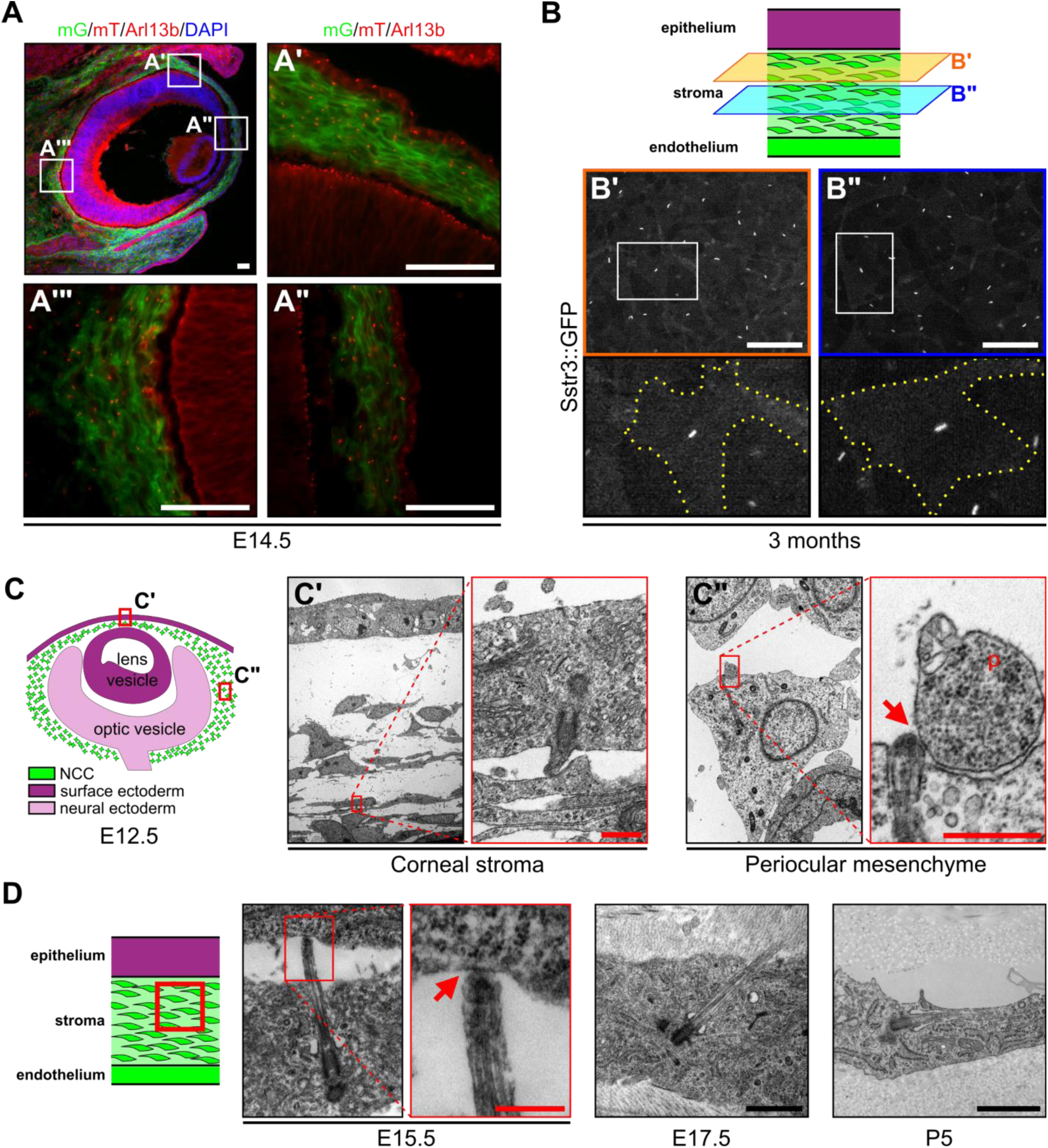
NCC of the periocular mesenchyme are ciliated. (**A**) Representative eye section of a *Wnt1-Cre;mT/mG* embryo at E14.5. NCC express the mG reporter (green cells) whereas cells from other embryonic origin express the mT reporter (red cells). Primary cilia were stained with an anti-Arl13b Ab and appear as small red rods. Scale bar, 50 μm. (**B**) Representative corneal stroma pictures of an Sstr3::GFP mouse at 3 months, in which primary cilia are GFP fluorescent. All stroma keratocytes are ciliated at adulthood. Scale bar, 50 μm. (**C**) Representative pictures of primary cilia in the corneal stroma and the periocular mesenchyme at E12.5. Scale bar, 0.5 μm. (**D**) Representative pictures of primary cilia in the corneal stroma at E15.5, E17.5, and P5. Scale bar, 1 μm. Primary cilia interact with neighboring cells or their cytoplasmic protrusions (red arrows). p, cytoplasmic protrusion.

**Figure 2:**
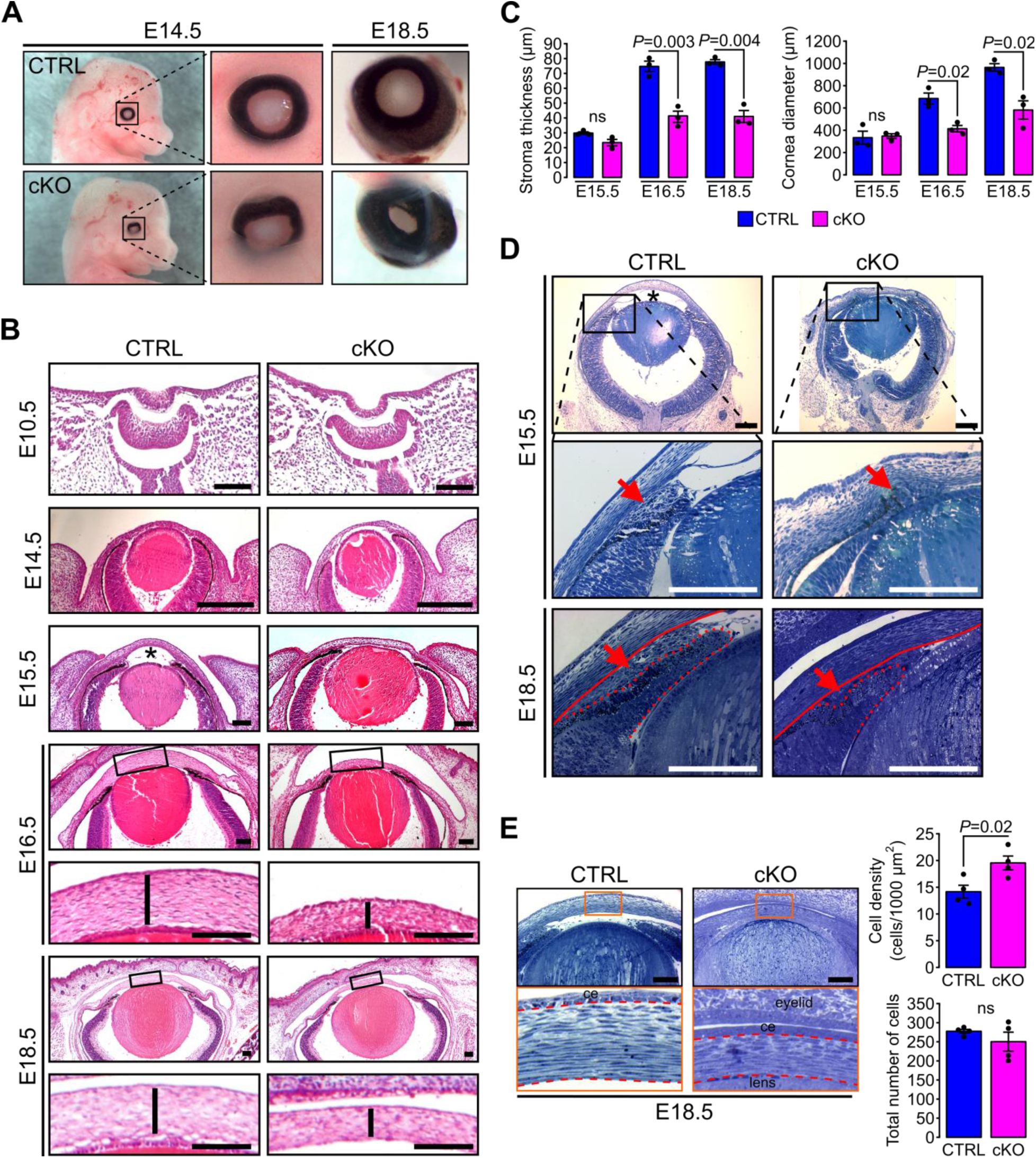
Primary cilium ablation in NCC leads to ASD phenotype. (**A**) Representative pictures of control and cKO eyes at E14.5 and E18.5. Enlarged pictures show the misorientation of the eyeball in the cKO embryos at E14.5 and the reduced size of the cornea at E18.5. (**B**) Representative eye sections stained with HE of control and cKO embryos at E10.5, E14.5, E15.5, E16.5, and E18.5. Boxed regions indicate the areas shown at higher magnification below. Scale bar, 100 μm. (**C**) The corneal stroma thickness and the corneal diameter were measured in the center of the cornea on HE stained paraffin sections at E15.5, E16.5, and E18.5 (n=3 embryos/group). (**D**) Representative plastic sections of the iridocorneal angle of control and cKO embryos at E15.5 and E18.5. Red arrows show the accumulation of mesenchymal cells between the cornea (underlined by the red line) and the presumptive iris (surrounded by the red dotted line). Scale bar, 150 μm; *, anterior chamber. (**E**) Representative plastic sections of corneas of control and cKO embryos at E18.5. Boxed regions indicate the areas shown at higher magnification below. Keratocytes were counted in the stroma which is surrounded by red dotted lines. The keratocyte density is significantly increased in cKO embryos compared to control but not the total number of cells (n=4 embryos/group). Scale bar, 100 μm; ce, corneal epithelium. Data are presented as mean SEM. Statistical significance was assessed using two-tailed Student’s *t*-test. ns, non-significant, P ≥ 0.05.

To further characterize the mutant phenotype of the AS, we conducted histological analysis of paraffin and plastic embedded samples (**Figure 2**). The eye field of the cKO embryos at E10.5 and 14.5 appeared indistinguishable from that of the control (**Figure 2B**). In contrast, at E15.5, we observed a significant reduction of the anterior chamber in the cKO (**Figure 2B-D**). At this stage, the mutant was also lacking most of the mesenchymal cells condensing at the developing iridocorneal angle between the cornea and the presumptive iris that was instead clearly visible in the control (**Figure 2D**). As a result, the iridocorneal area appeared disorganized in the mutant with a narrower angle than in the control. Between E16.5 and E18.5, the corneal stroma thickness and the corneal diameter were both significantly reduced in the cKO embryo compared to the control (**Figure 2B-C**). Moreover, the iridocorneal angle abnormalities found in the mutant eye persisted with the presumptive iris significantly shorter than that of the control (**Figure 2D**). Plastic cross sections of the cornea also revealed that the density of the keratocytes was significantly higher in the cKO stroma compared to control however, the total number of cells in corneal stroma remained unchanged in both genotypes (**Figure 2E**). Thus, the increased keratocyte density detected in the cKO embryos was essentially due to its smaller volume cornea compared to that of control. Moreover, the unchanged number of keratocytes in the stroma of both genotypes suggests that the primary cilium ablation did not impair the NCC migration into the corneal stroma. Thus, the ablation of the primary cilium in the NCC leads to ASD that is not due to NCC migration defects.

Next, we have examined the tridimensional organization and the morphology of keratocytes. Keratocyte morphology and spatial distribution across the stroma vary in mammals, with an increased density, flattening, and extension of the keratocytes in the posterior part of the stroma in late embryonic development [50, 51]. To determine stromal organization, we developed a quantitative tool using live confocal imaging on mT/mG mice. Segmentation and quantification of the stromal extracellular spaces was carried out with ilastik [52] (**Supplement Figure 2**). We determined the amount (%) and the average size of segmented extracellular spaces using Fiji [53]. In control embryos, the amount of extracellular spaces was significantly lower in the posterior than the anterior portion of the corneal stroma, defining an antero-posterior gradient of the keratocyte density (**Figure 3B**). Moreover, the size of the extracellular spaces in the posterior part was significantly smaller than in the anterior part of the stroma in control embryos, defining an antero-posterior gradient of the extracellular space size (**Figure 3C**). In cKO embryos, the antero-posterior gradient of the keratocyte density was maintained. However, that of the average size of extracellular spaces was not (**Figure 3B**). In addition, the amount of extracellular spaces was significantly reduced in both the anterior and posterior parts of the cKO stroma compared to control (**Figure 3B-C**). Thus, the keratocyte distribution in the corneal stroma of the cKO embryos was denser than that of the control. Moreover, the differences in the anterior-posterior gradient of the amount of extracellular space and the average size of single extracellular spaces are abnormal in the cKO suggesting a defective tridimensional organization of the keratocytes in the mutant. Thus, the ablation of the primary cilium in NCC impairs the spatial organization of keratocytes in the stroma independently from cell migration.

**Figure 3:**
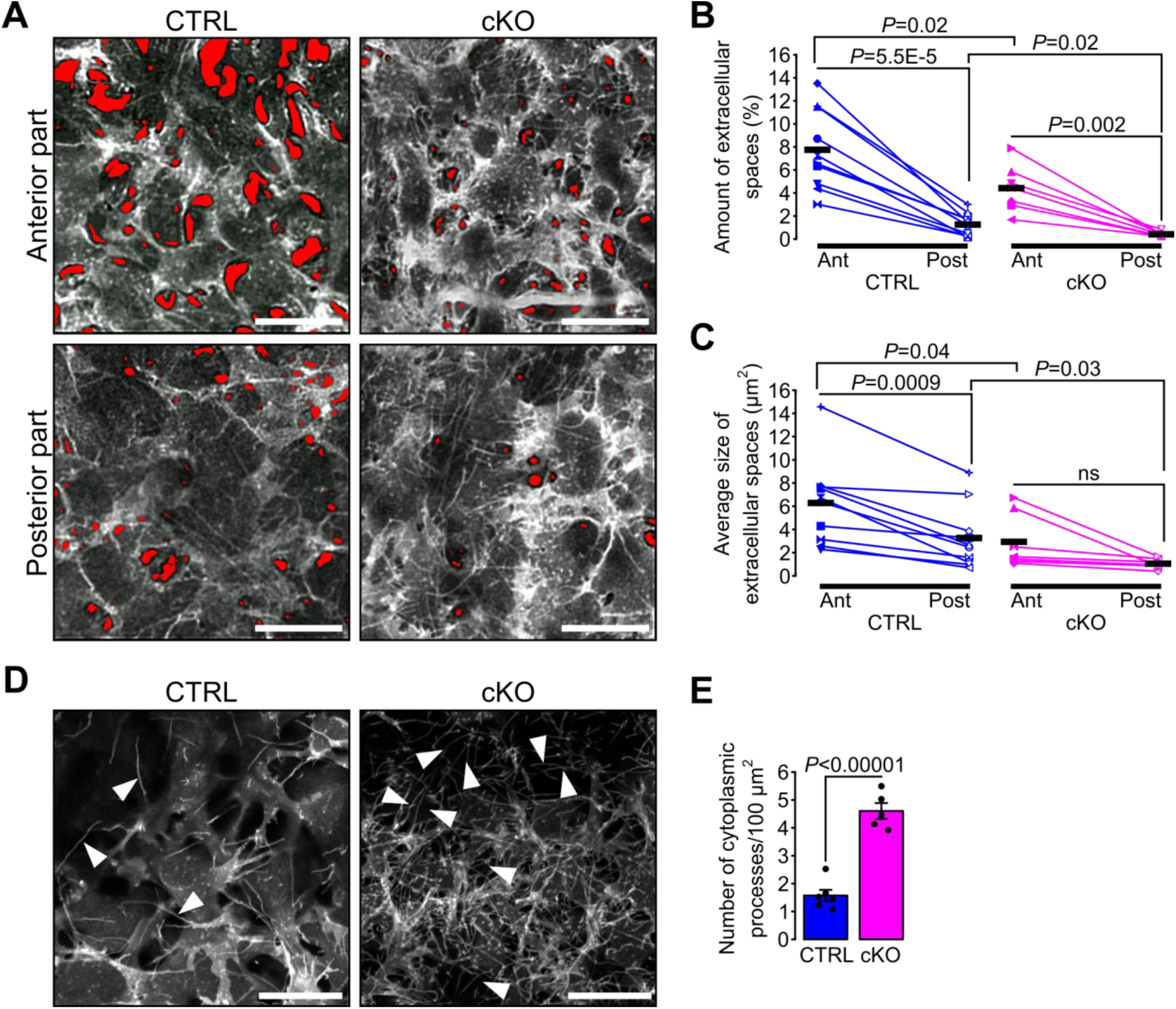
Primary cilium ablation in NCC impairs the spatial organization and the morphology of keratocytes. (**A**) Representative live confocal pictures at different levels of the corneal stroma of E18.5 cKO and control embryos crossed with the mT/mG reporter line which allows direct visualization of the cell membrane without any staining. Here, only the mG fluorescent signal is displayed. The red areas are the extracellular spaces segmented with Ilastik. Pictures of the anterior and posterior parts were picked from the first tenth of the stroma underlying the corneal epithelium and from the last tenth of the stroma just on top of the corneal endothelium, respectively. Scale bar, 25 μm. (**B-C**) The amount and the average size of extracellular spaces were determined in the anterior (Ant) and the posterior (Post) parts of the corneal stroma. The amount of extracellular spaces corresponds to the sum of the area of the extracellular spaces normalized to the total area of the picture. Each symbol represents the mean value of an embryo obtained from 5-12 pictures. Black lines represent the mean value of each group (n=10 control, n=7 cKO). Statistical significance was assessed using two-tailed Student’s *t*-test between the two genotypes, and using paired two-tailed Student’s *t*-test between the anterior and posterior side measurements from mice with the same genotype. ns, non-significant, P ≥ 0.05. (**D**) Representative confocal pictures of the first layer of stromal keratocytes underlying the corneal epithelium (only the mG reporter is displayed). Cytoplasmic processes (white arrowheads) are more numerous in the cKO embryos. Scale bar, 20 μm. (**E**) Quantification of the number of cytoplasmic processes in the first layer of keratocytes underlying the corneal epithelium (n=5-6 embryos/group). Data are presented as mean SEM. Statistical significance was assessed using two-tailed Student’s *t*-test.

Because the density and the spatial organization of the keratocytes are abnormal in the mutant stroma we sought to investigate the morphology of single keratocytes *in vivo*. Keratocytes are characterized by their cytoplasmic processes interconnected with each other to form a dense and complex 3D network [54]. We focused on the junction between the corneal epithelium and the stroma because here the density of keratocytes is lowest. In both genotypes, we could distinguish single cytoplasmic processes of the first keratocyte layer expressing the mG reporter underneath the overlying corneal epithelial cells layer expressing the mT reporter. The number of cytoplasmic processes was significantly increased in the cKO embryos compared to control (**Figure 3D-E**), suggesting a possible role of the primary cilium in controlling the morphology and the number of cytoplasmic processes of keratocytes.

### Ablation of Ift88 in NCC disrupts Hh signaling in a subpopulation of POM cells surrounding RPE and at the iridocorneal angle

Morphological changes, as well as spatial organization of the cells in a tissue, occur as cells differentiate. This implies a wide variety of signaling pathways occurring in a timed and coordinated fashion. Mice lacking heparan sulfate in NCC display ASD phenotypes resembling those caused by the lack of NCC cilia including cornea stroma hypoplasia, dysgenesis of the iridocorneal angle and decreased depth of the anterior chamber [55]. Because in these mice the TGFβ2 pathway is disrupted, we assessed whether the primary cilium ablation in the NCC affected the TGFβ2 pathway in the cornea. At E18.5, the expression of *Tgfbr1*, *Tgfbr2*, *Smad2*, *Smad3*, *Smad4* and *Smad7* and the percentage of pSmad2/3^+^ cells in the cornea remained indistinguishable between the cKO and control mice (**Supplement Figure 3**).

Hh, Wnt and Notch signaling pathways have been linked to the primary cilium [22]. We therefore used quantitative real-time PCR (RT-qPCR) on isolated ocular NCC and reporter mouse lines to examine the expression of specific target genes of the above-mentioned pathways. To isolate ocular NCC, cKO and control mice were crossed to the mT/mG transgenic reporter line (**Supplement Figure 1**) and mG^+^ cells of the dissected sclera and cornea tissue were digested and FACS sorted following the approach shown in (**Figure 4A**). NCC from the E18.5 cKO mouse showed a significant decrease of Hh target genes including *Gli1* and *Ptch1* but not *CyclinD* when compared to control NCC (**Figure 4B**). In contrast, the expression of target genes of the Wnt and Notch pathways, including *Axin2*, *β-catenin*, *Lef1*, *Hey1*, *Hes1* and *Maml1* remained unchanged between mutant and control (**Figure 4B**).

**Figure 4:**
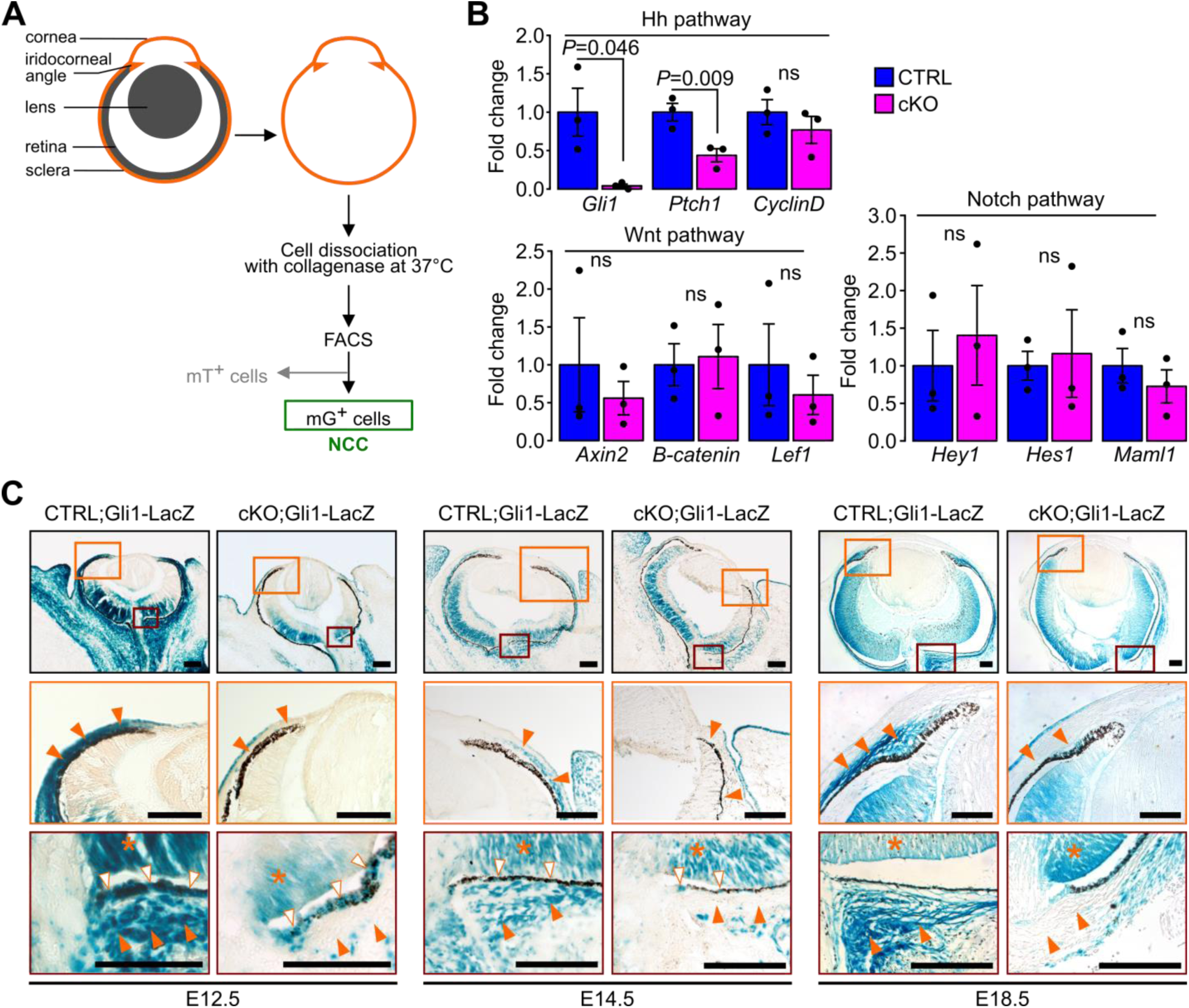
Ablation of Ift88 in NCC reduces Hh signaling in the POM NCC. (**A**) Workflow followed to collect the NCC from the eyeball by cell sorting. (**B**) Hh, Wnt and Notch target gene expression in NCC at E18.5. Fold change is expressed as mean ± SEM (n=3 embryos/group, both eyes of each embryo were pooled together). Only the expression of Hh pathway target genes was affected by the primary cilium ablation in the NCC. Statistical significance was assessed using two-tailed Student’s *t*-test. ns, non-significant, P ≥ 0.05. (**C**) Hh activity assessed by Gli1-LacZ staining throughout the embryonic development. In the control embryos, the Hh activity progressively decreases in the POM from E12.5 to E18.5 but remains activated in a subpopulation of POM cells surrounding the RPE layer and extending until the iridocorneal angle, as well as in the POM surrounding the optic nerve (orange arrowheads). The primary cilium ablation in NCC leads to the absence of Hh activity in these specific areas. Hh signaling remains active in non-NCC derived tissues in the cKO embryos like in the RPE (white arrowheads) and the retina (asterisks). Scale bar, 100 μm.

To determine the spatial distribution of Hh responsive cells of the POM during eye development we crossed cKO and control mice with the *Gli1-LacZ* reporter mouse line [56]. In control mice at E12.5, we detected intense Hh activity in most of the POM, given that all the cells were strongly LacZ-positive (**Figure 4C**). At E14.5 and E18.5, the Hh activity in the POM was greatly decreased with only a subpopulation of POM cells surrounding the RPE layer remaining positive. This layer of Hh responsive cells extended from the optic nerve area to the iridocorneal angle in control embryos (**Figure 4C**). By adulthood, the Hh signaling remained active only in the choroid area, around the optic nerve, and in the ciliary body (**Supplement Figure 4**). In contrast, in cKO mice, the Hh activity was nearly completely absent in the POM at any developmental stages. In both mutant and control mice, we detected intense Hh activity in tissues derived from the neuroectoderm including the retina and RPE at all developmental stages (**Figure 4C**) [57]. Interestingly, no Hh activity was detected in the cornea of both cKO and mutant mice between E12.5 and E18.5 despite the presence of cilia in keratocyte precursors throughout development (**Figure 4C**). Thus, ablation of the primary cilium in NCC disrupted the Hh activity in POM.

The Hh signaling pathway plays an important role in promoting cell proliferation in different tissues and cell types [58]. Thus, we hypothesized that the Hh activity could influence cell proliferation rates in the Hh-active POM subpopulation surrounding the RPE cell layer. We tested this possibility in E14.5 embryos. At this developmental stage, the Hh activity in the POM was restricted to a 20 µm thick cell layer (0-20 µm) as measured from the RPE cell layer in sagittal sections of the eye (**Figure 5A**). In addition, we also arbitrarily defined a Hh-negative area of the POM encompassed between 20 µm and 50 μm from the RPE cell layer (20-50 µm) functioning as an internal control together with the Hh-negative cornea. Proliferation rates were assessed by counting the number of cells stained with an anti-Ki67 Ab and normalized to the total number of cells in the same area stained with DAPI. At E14.5, cell proliferation was significantly decreased in the POM close to the RPE layer (0-20µm) of the cKO mutants compared to control (**Figure 5B**). In contrast, proliferation rates remained unchanged in the outer region of the POM (20-50 µm) and in the cornea where the Hh was normally not activated (**Figure 5B**). Thus, ablation of the primary cilium in NCC leads to a decreased cell proliferation specifically in a subpopulation of POM cell surrounding the RPE due to local inactivation of the Hh signaling.

**Figure 5:**
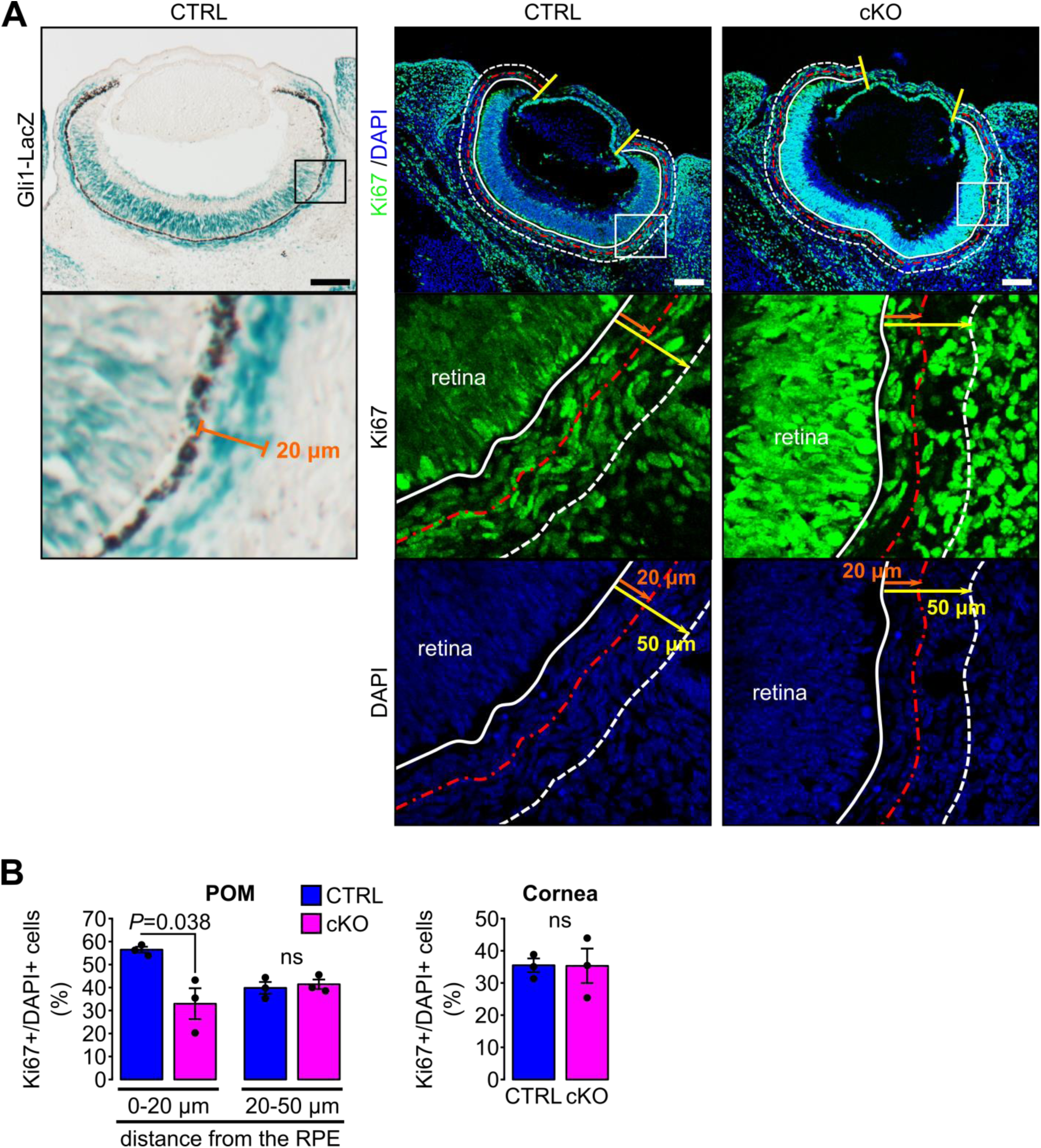
Primary cilium ablation in NCC decreases the cell proliferation specifically in the POM cells surrounding the RPE layer. (**A**) Cell proliferation was assessed at E14.5 in the cornea (surrounded by yellow lines) and in the POM. The white line indicates the RPE layer. Two areas were defined in the POM: the first one between 0 and 20 μm from the RPE layer (delineated with the red dotted line), corresponding to the mean thickness of Gli1LacZ-positive POM cells surrounding the RPE, and the second one from 20 to 50 μm from the RPE layer (delineated with the white dotted line). Scale bar, 100 μm. (**B**) Percentage of Ki67^+^ cells in the different areas of the POM and in the cornea. The number of Ki67^+^/BrdU^+^ cells is expressed as mean ± SEM. Statistical significance was assessed using one-tailed Student’s *t*-test (n=3 embryos/group). ns, non-significant, P ≥ 0.05.

### Embryonic POM cells of the sclera progressively lose their ability to respond to Ihh produced by choroidal endothelial cells

Next, we sought to determine how the Hh signaling is maintained in a restricted subpopulation of NCC within the POM. Because mammals express three different Hh homologues, *Dhh*, *Ihh* and *Shh*, we analyzed dissected tissue of the eye from E18.5 mice, including NCC by RT-qPCR to determine which Hh ligand is expressed in NCC derived tissue of the eye as shown in the scheme in **Figure 6A**. We found that the expression of *Ihh* was significantly higher than that of *Dhh* and *Shh* in the sclera including the RPE layer but not in the cornea (**Figure 6A**). As control, we found that *Shh* but not *Dhh* and *Ihh* was expressed in the isolated neural retina where NCC are excluded (**Figure 6A**).

**Figure 6:**
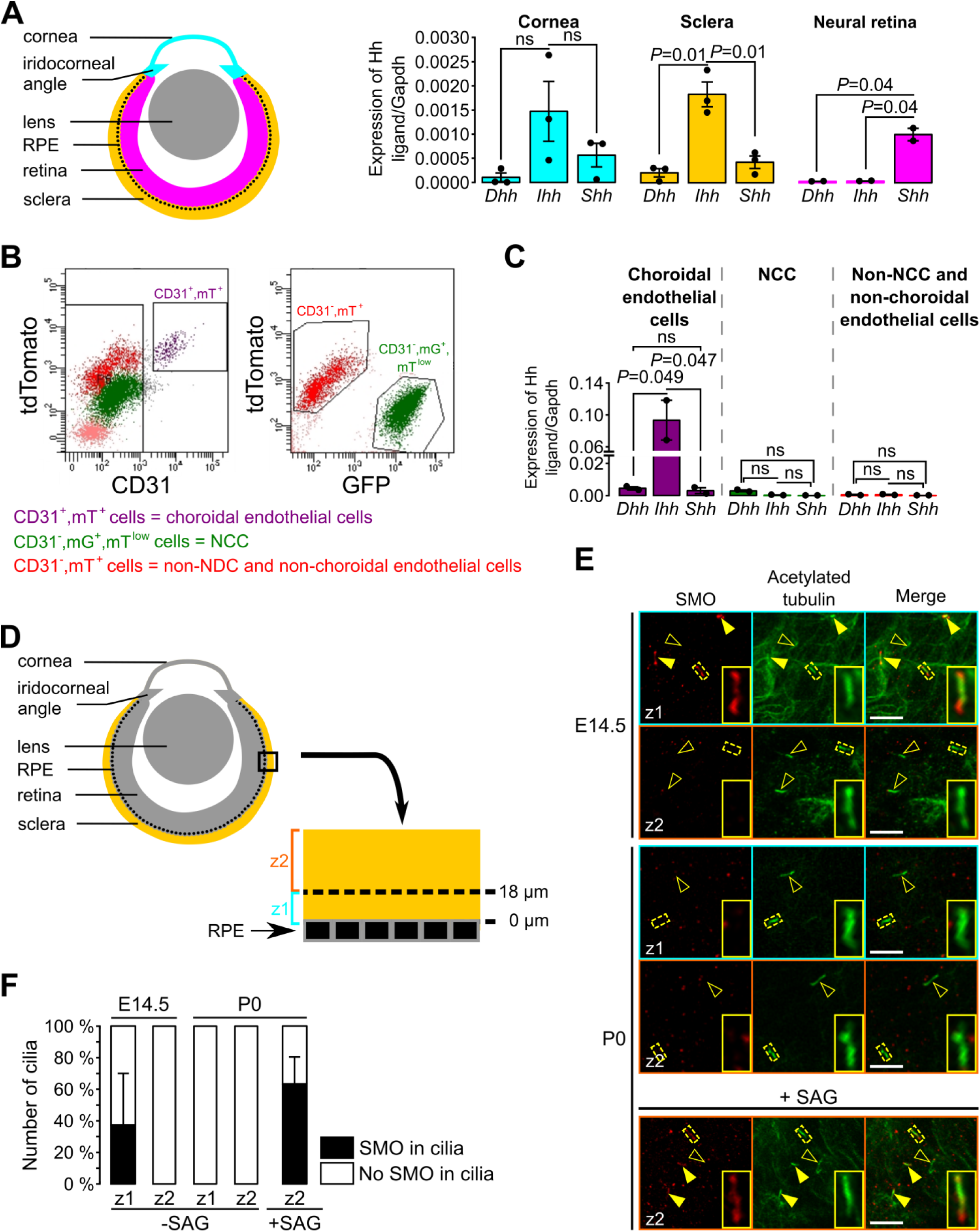
During the embryonic development, POM cells of the sclera lose their ability to respond to Ihh produced by choroidal endothelial cells. (**A**) Expression of *Dhh*, *Ihh* and *Shh* in the cornea (including the iridocorneal angle), the neural retina and the posterior half of the eyeball without the neural retina (corresponding to the sclera, choroid and RPE layer, here indicated as the sclera) in control embryos at E18.5 (n=2-3). The different colors correspond to the different dissected parts of the eyeball as shown on the schematic representation on the left. (**B**) Representative bivariate dot plots of isolated cells from E18.5 embryonic eyeballs gated on mT and CD31 or mG expression. (**C**) Expression of *Dhh*, *Ihh,* and *Shh* in sorted cells isolated from E18.5 embryonic eyeballs (N=2 independent experiments; n=4-6 eyeballs/N). Results are mean ± SEM. Statistical significance was assessed using one-way ANOVA with posthoc Tukey HSD test. ns, non-significant, P ≥ 0.05. (**D**) Schematic representation of the areas imaged in the sclera by confocal microscopy. On E14.5 embryonic sclera, we identified two different layers: z1 from 0 and 18 μm from the RPE in which SMO is accumulated in primary cilia, and z2 above 18 μm from the RPE in which SMO is not accumulated in primary cilia. For further quantifications at P0, the same layers were considered. (**E**) Representative pictures of SMO and acetylated tubulin (primary cilia) staining extracted from z1 and z2 z-stacks in whole-mount scleras at E14.5 and P0. Primary cilia with SMO accumulation are indicated with yellow arrowheads, whereas primary cilia without SMO are indicated by empty arrowheads. At P0, SMO is visible in primary cilia only upon SAG stimulation. Scale bar, 5 μm. (**F**) Quantification of the number of primary cilia in which SMO is present or absent (n=5 at E14.5, n=3 at P0).

To determine the source of the Hh ligand we sorted and analyzed by RT-qPCR different cell types of the sclera. A previous study showed by *in situ* hybridization and immunohistochemistry an overlapping staining between *Ihh* and collagen type IV, a vascular endothelial marker, suggesting a possible involvement of the choroid endothelium in *Ihh* production [59]. Thus, we isolated by FACS choroidal endothelial cells using the mT/mG fluorescent reporter combined to an anti-CD31 Ab (a specific marker of endothelial cells). Three different cell populations were sorted: choroidal endothelial cells (CD31^+^, mT^+^, mG^-^), NCC (CD31^-^, mT^low^, mG^+^) and cells which are neither NCC nor choroidal endothelial cells (CD31^-^, mT^+^, mG^-^; **Figure 6B**). At E18.5, among the non-NCC, *Ihh* was significantly more expressed by the choroidal endothelial cells than *Dhh* and *Shh* (**Figure 6C**). Thus, we provided molecular evidence that endothelial cells of the POM produce the Hh ligand *Ihh* in the sclera.

As shown in **Figure 4** during eye development, NCC of the peripheral POM progressively lost the Hh signaling activity while Hh-responsive cells remained confined to the choroid area and at the iridocorneal angle. To gain mechanistic understanding on how this process occurs we analyzed the presence of cilia and Hh components in the ciliary compartment of Hh-negative POM cells. In presence of an Hh ligand, its receptor PTCH exits the ciliary compartment allowing SMO to concentrate into the cilium, which is an essential step to activate the Hh signaling [25]. To determine whether POM cells were still ciliated and able to respond to a Hh signal, we stained primary cilia and SMO in whole mount preparations of the wild-type sclera. We collected confocal optical sections of the POM between the RPE layer and the periphery of the sclera as indicated in (**Figure 6D**). At E14.5 and P0, all cells of the POM were ciliated (**Figure 6E**). Thus, we excluded the possibility that NCC of the peripheral POM lost the Hh activity due to resorption of the primary cilium. At E14.5, we observed an accumulation of SMO in primary cilia of the inner cell layers of the POM surrounding the RPE (0-18 μm) consistent with the Hh activity detected in these cell layers (**Figure 4**). In contrast, we did not detect SMO in the primary cilium in cells of the peripheral POM (>18 μm from the RPE) (**Figure 6E-F**). At P0, SMO was undetectable in primary cilia of NCC of the entire POM (**Figure 6E-F**). However, when eyeballs from P0 mice were treated with SAG, SMO accumulated in cilia of POM cells (**Figure 6E-F**). This implies that POM cells of the P0 sclera were still able to activate Hh signaling upon ligand stimulation, and suggests that the decreased Hh activity in the POM during ocular development is controlled by the diffusion in the POM of the Hh ligand *Ihh*.

### Primary cilium ablation in NCC leads to corneal neovascularization

Heterozygote mutations in *FOXC1* and *PITX2* genes account for ∼40% of ASD cases [60]. In mice, *Foxc1* and *Pitx2* haploinsufficiency leads to ocular phenotypes recapitulating human conditions of ASD including iridocorneal angle abnormalities, thinning and abnormal vascularization of the cornea [61–64]. Because these phenotypes were also detected in the cKO mouse, we tested whether absence of cilia in the NCC affected the expression of ASD genes in E18.5 cKO embryonic corneas. RT-qPCR revealed that gene expression of *Foxc1* and *Pitx2* was significantly decreased in corneas of cKO embryos compared to controls while *Pax6* expression remained unchanged (**Figure 7A**). Because *FOXC1* and *PITX2* are indispensable to specify corneal angiogenic privilege [1, 62–64] we examined the neovascularization process in the cKO mutant. To visualize blood vessels, we stained whole corneas with an Ab directed to endomucin, a marker of the vascular endothelium. Abnormal spread of blood vessels was detected into the corneas of E18.5 cKO mutant embryos (**Figure 7B-C**) where vessels covered about 8% of the total corneal area (**Figure 7D**). To gain additional insights on the abnormal neovascularization process, we performed live imaging using confocal microscopy at the cornea periphery using enucleated eyes from E18.5 embryos with the *mT/mG* reporter. Due to their mesenchymal origin, blood vessels were easily detectable as red (mT) tubular network clearly distinct from the green (mG) NCC-derived POM (**Supplement Figure 5**). By confocal optical sectioning we identified the major arterial circle which served as reproducible reference between the end of the sclera and the beginning of the cornea. The average area occupied by blood vessels in microscope fields selected at the corneal periphery, occupied ∼30% of the total area in the cKO but only ∼4% in the control (**Figure 7E-F**). In addition, while peripheral blood vessels of the control were present only in the superficial layers of the sclera-cornea interface on both sides of the major arterial circle (∼20% of the thickness), those of the mutant invaded the lower layers of both, the sclera (∼50% of the thickness) and the cornea (∼80% of the thickness). These results demonstrate that ablation of the primary cilium in the NCC leads to the loss of the angiogenic privilege in the cornea as well as vascular abnormalities in the sclera, potentially due to the lowered expression of *Foxc1* and *Pitx2*.

**Figure 7:**
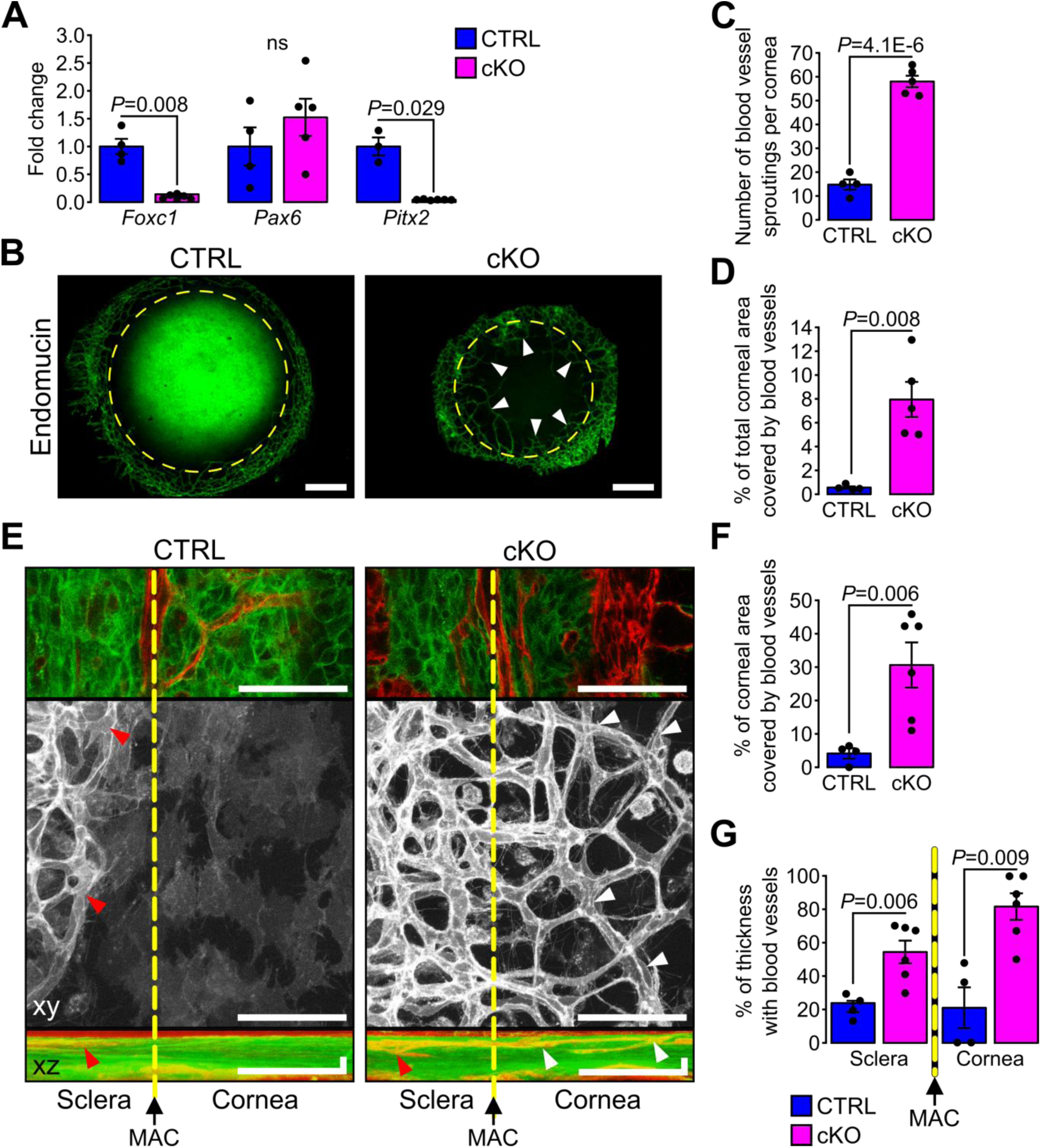
Primary cilium ablation in NCC leads to corneal neovascularization. (**A**) *Foxc1*, *Pax6* and *Pitx2* expression in the cornea (including the iridocorneal angle) (n=3-6 embryos/group). (**B**) Representative pictures of whole-mount corneas stained with an anti-endomucin Ab in control and cKO embryos at E18.5. Blood vessels are indicated by white arrowheads and the yellow dotted line surrounds the cornea. Scale bar, 250 μm. (**C-D**) Quantification of the number of blood vessel sproutings and the total corneal area covered by blood vessels in E18.5 control and cKO embryos (n=4-5 embryos/group). We defined the vessel point crossing the corneal border and extending forward to the cornea as vessel sprouting. (**E**) Representative confocal pictures of the cornea periphery at E18.5. The yellow dotted line represents the corneal boundary, established by the position of the major arterial circle (MAC, picture on the top). Only the mT reporter is displayed on the xy maximum intensity projection (grayscale picture), whereas both mT and mG reporters are displayed on the xz maximum intensity projection. Arrowheads indicate blood vessels which are only visible in the sclera side in controls (red arrowheads) whereas they extend into the cornea in the cKO embryos (white arrowheads). Scale bar, 25 μm. (**F**) Percentage of the corneal area covered by blood vessels at the periphery of the cornea (n=4-6 embryos/group). (**G**) Quantification of the corneal and stroma thickness in which blood vessels are present (n=4-6 embryos/group). Results are mean ± SD. Statistical significance was assessed using two-tailed Student’s *t*-test. ns, non-significant, P ≥ 0.05.

### Primary cilium ablation in NCC impairs early corneal innervation and centripetal migration of the sensory nerves

Concurrent with the establishment of the angiogenic privilege is corneal innervation [65]. Because corneal sensory nerves are derived from the neural crest that is part of the trigeminal ganglion [66], we assessed whether primary cilium ablation in NCC would also impact the corneal innervation. During development, corneas of both control and cKO embryos were innervated (**Figure 8A**). However, at E13.5, the number of nerve bundles detected in the cKO eye was significantly lower compared to that of the control (**Figure 8B**). Moreover, abnormally long nerve projections across the cornea were observed in cKO embryos whereas nerves are only visible at the periphery of the cornea in control (**Figure 8A**). At E18.5, corneas of both genotypes were fully innervated (**Figure 8A**), including the corneal epithelium (**Supplement Figure 6**). However, the corneal nerve density was significantly higher in the cKO embryos (**Figure 8C**). In addition, the nerves in cKO corneas were less organized centripetally compared to the control. The average angle θ formed by the major nerve branches with the radius of the cornea was significantly bigger in cKO embryos compared to controls (**Figure 8D-E**). These data suggested that the primary cilium ablation in the NCC impaired the early corneal innervation and the centripetal migration of the sensory nerves.

**Figure 8:**
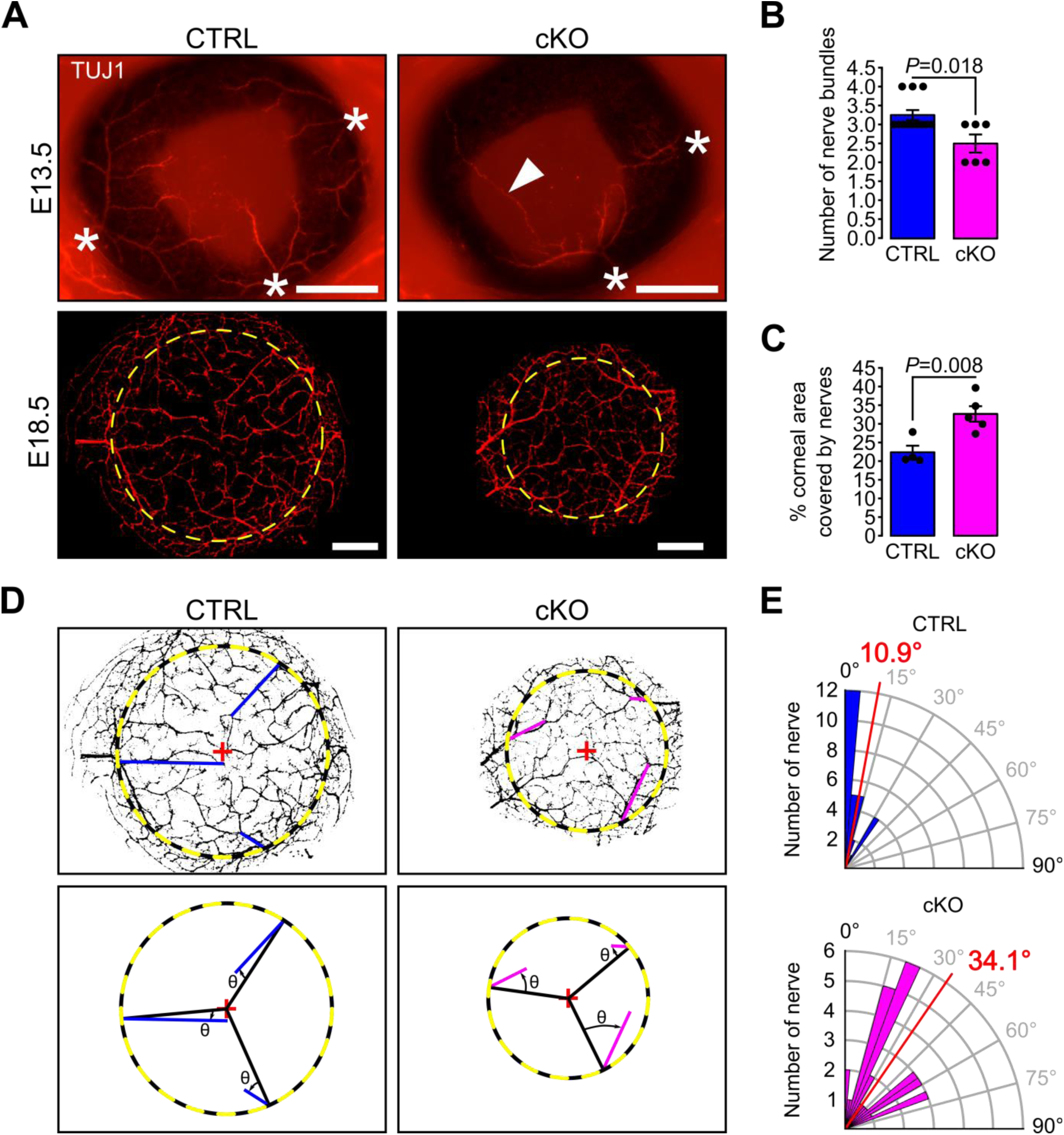
Primary cilium ablation in NCC reduces early corneal innervation and impairs the centripetal migration of sensory nerves. (**A**) Representative pictures of whole-mount corneas stained with an anti-TUJ1 Ab in control and cKO embryos at E13.5 and E18.5. At E13.5, three to four nerves bundles (asterisk) are present in controls and start to project at the periphery of the cornea. Abnormal nerve projections (head arrow) across the cornea are visible in some cKO embryos at E13.5. By E18.5, the cornea (surrounded by the yellow dotted line) is fully innervated in both the control and cKO embryos. Scale bar, 250 μm. (**B**) Quantification of the number of nerve bundles at E13.5 which is significantly decreased in cKO embryos compared to control (n=8-13 embryos/group). (**C**) Nerve density at E18.5 quantified by measuring the percentage of corneal area covered by nerves (n=4-5 embryos/group). (**D**) Corneal nerve segmentation from E18.5 embryos. The colored lines (blue in control, pink in cKO) represent the axis of major nerve branches formed between the point where the main nerve branch crosses the corneal border (yellow dotted line) and its end point (final point of the nerve branch or the branching point from which two secondary branches with similar diameter arise). The angle θ formed by the major nerve branches and the radius of the cornea (black line) was measured with Fiji. (**E**) Quantification of the angle θ in control and cKO mice at E18.5. The nerves in the cKO embryos are less centripetally organized than in control (*P* = 8.43E^-7^, n=4-5 embryos/group). In B and C, results are mean ± SEM. In D, the mean angle is indicated in red. Statistical significance was assessed using two-tailed Student’s *t*-test.

## DISCUSSION

Ocular conditions such as microphthalmos/anophthalmos, aniridia, sclerocornea, abnormal corneal thickness, corneal neovascularization, abnormal iridocorneal angle, corneal opacity, glaucoma, and cataract, have been reported in patients affected by ciliopaties including Joubert and Mekel syndrome, suggesting an implication of the primary cilium in AS development [27–32]. However, the pathogenesis of these conditions remains largely unknown. In the present study, we have demonstrated that conditional ablation of the primary cilium in NCC led to ASD with conditions similar to those observed in humans. We showed that primary cilia are present in virtually all neural crest-derived cells of the POM including keratocytes. However, we found that primary cilia are required for the propagation of the Hh pathway in a subpopulation of POM cells surrounding the RPE layer but not in neural crest-derived cells of the cornea where the Hh pathway is not active. We showed that the expression of ASD-causing genes including *Foxc1* and *Pitx2* was strongly reduced in the cornea and iridocorneal angle tissue lacking cilia in the NCC which is consistent with AS phenotypes, including the loss of corneal angiogenic privilege. Thus, we have established an important functional connection between cilia-dependent signaling and ASD. Overall, this study demonstrated the novel and fundamental role of the primary cilium in the development of structures of the AS derived from the NCC.

Mice carrying somatic null mutations in genes encoding proteins required for cilia assembly or ciliary function display severe microphthalmia and anophthalmia [38, 43, 67–71]. This conditions are likely due to aberrant Hh activity in cells of the neuroepithelium of the developing optic vesicle that precedes the development of the AS [72]. To overcome this obstacle and test the function of the primary cilium in AS development we have ablated IFT88 in the NCC, a multipotent progenitor cell population that produces a large range of differentiation fates including a large portion of AS structures [5]. Relevant to our study, conditional deletions of ASD causative genes such as *Foxc1*, *Pitx2* or *Ext1* in NCC, lead to defects of the AS including anophthalmos, abnormal corneal thickness, reduced anterior chamber, defective iridocorneal angle, and corneal neovascularization similar to abnormalities observed in ASD patients carring dominant mutations in these genes [55, 64, 73]. However, although the cKO mice revealed a direct connection between the primary cilium and ASD, its severe craniofacial defects caused late embryonic death and prevented us to follow the postnatal development of structures such as the iris, trabecular meshwork and Schlemm’s canal [8]. Thus, further studies involving conditional ablation of the cilium in specific AS tissues will be needful.

The low expression levels of *Foxc1* and *Pitx2* detected in the cornea of the cKO mutant could be at the origin of some of its AS phenotypes, including abnormal corneal vascularzation and thinning of the cornea [1, 60–64, 73, 74]. Mice carrying only one copy of *Foxc1* (*NC-Foxc1^+/-^*) in the NCC do not lose cornea angiogenic privilege so their cornea remains avascular. Whereas loss of both *Foxc1* copies in the NCC leads to severe neovascularization of the whole cornea suggesting a strong relationship between the maintenance of the corneal angiogenic privilege and the dosage of *Foxc1* [64]. Consistent with this possibility, we observed a partial vascularization in the cornea of the cKO mouse. The dosage of *Foxc1* and *Pitx2* also regulates differentially the corneal thickness. While the corneal thickness increases in *NC-Foxc1^-/-^* mice and *Pitx2^-/-^*, it remains unaltered in *NC-Foxc1^+/-^* and decreases in *Pitx2^+/-^*mice, respectively [61, 64, 75]. Thus, the reduction of *Pitx2* expression could be one of the concomitant causes leading to thinning of the cornea observed in the cKO embryos. In summary, the partial corneal vascularization and the decreased stroma thickness observed in the cKO embryos are coherent with a decreased expression of *Foxc1* and *Pitx2* in the cornea.

The abnormal corneal neovascularization observed in the cKO could also derive from the reduction or absence of the Hh activity in the POM. In a zebrafish model, the loss of Hh signaling induces excess sprouting of the blood vessels in the dorsal eye and impaires growth of the blood vessels in the ventral eye [76]. These observations in the eye tissue of the fish are also consistent with the aberrant expansion of the blood vessels domain deep into the sclera of cKO embryos (**Figure 7**). A thorough analysis of the choroid in NCC conditional mutants with a reduced Hh activity obtained independently from the ablation of the primary cilium could elucidate a specific role of the Hh pathway in development and patterning of the choroid.

The decreased expression of *Foxc1* and *Pitx2* in tissues of the AS of the cKO mouse suggests a possible direct or indirect mechanism of gene expression regulation of these transcription factors by cilia-dependent signaling. A recent study has shown that mutations in the *FOXC1* gene in humans cause, in addition to ASD, systemic conditions similar to those found in ciliopathic patients including polydactyly (ARVO meeting 2019 abstract #1617, personal communication). This study also shows that lowering the level of *Foxc1* expression in epithelial cells induces a reduction of cilia length and interferes with cilia localization of Hh components, notably GLI2 (ARVO meeting 2019 abstract #1617, personal communication). FOXC1 also appears to directly interact with GLI2 and promote the expression of Ihh target genes, including *Gli1* and *Ptch1* [77]. Together, these data suggest a strong relation between primary cilia, *Foxc1* expression and Hh activation. This could explain why similar ocular and systemic phenotypes are observed in ASD and ciliopathic patients.

The process of corneal innervation occurs in parallel with the establishment of the corneal angiogenic privilege. The majority of corneal nerves derive from the ophthalmic division of the trigeminal ganglion of NCC origin [66]. The cKO mouse displays a defective corneal innervation patterning consisting in abnormalities of the nerve projections similar to those described in a *Semaphorin 3* deficient mouse models [78, 79]. *Semaphorin 3* regulates the navigation process of the nerves by guiding the direction of axon growth [79]. In our ciliary model, the decreased number of nerve bundles at E13.5 and the less centripetal migration of the corneal nerves at E18.5 suggest that the navigation process of trigeminal nerves in the cornea is affected. Consistent with these exciting possibility, ciliogenic proteins including KIF3A, IFT88, and ARL13B, are required for the directional migration of GABAergic neurons [80, 81]. Thus, assembly of the primary cilium on trigeminal nerves could be critical in restablishment of the corneal innervation following corneal transplant or during recovery of trauma of the cornea.

The Hh signal transduction pathway is intimately linked to the primary cilium [23, 24], and its dosage is critical to ensure normal eye development [38, 82, 83]. In this study, we have shown that at an early stage of the AS development virtually all the POM cells are actively responding to Hh signaling. However, as the AS development progresses only a subpopulation of NCC of the POM sourrounding the RPE cell layer and in the iridocorneal angle, maintains Hh responsiveness by actively expressing *Gli1* and localizing SMO into the primary cilium compartment. Moreover, the NCC derived components of the cornea, which includes keratocyte precursors and corneal endothelium, were not found Hh responsive at any developmental stage. How is this differential distribution of Hh responsive achieved in cells of similar embryonic origin? Cells of the POM maintained the ability to respond to Hh stimulation as we have demonstrated at several levels: all cells of the POM and the corneal stroma were found ciliated thoughout development and after birth. In addition, *in vitro* SAG treatment of ocular tissues indicated that all POM cells were able to respond to Hh stimuli by promoting SMO localizing to their primary cilium. We have identified CD31 positive cells of the POM as the main source of Ihh. This suggests that endothelial cells of the choroid vasculature could be the principal source of Ihh in the eye. Consistent with this possibility, a previous study showed that *Ihh* expression in the POM colocalizes with collagen IV, a marker for the vasculature, next to the RPE layer [59]. Thus, the secretion of Ihh by the choroid vessels could regulate the propagation of the Hh responsivness in the periocular space and the cornea.

Our data suggest that one of the main functions of the Hh activity in the POM during late gestation is to promote cell division in the subpopulation of NCC sourrounding the RPE layer. Surprisingly, although the size of the cornea in the cKO mouse is considerably reduced, the number of keratocyte precursors within the corneal stroma is similar to the one found in control corneas. Thus, the reduction of cell proliferation in this subpopulation of POM cells does not account for the number of cells that populate the cornea. However, we have identified a group of mesenchymal cells at the iridocorneal angle of control mice that were absent or reduced in number in the cKO mutant. These cells could be precursors of drainage structures of the AS such as the trabecular meshwork and/or the iris stroma. Thus, comparative studies involving differential single cell analysis of the POM cells from the AS of the cKO and control mice will be crucial to identify this distinct cell population at the iridocorneal angle.

In conclusion, we have shown that primary cilia are indispensable for normal development of NCC-derived structures in the ocular AS. When the primary cilium is ablated in NCC, this leads to ASD phenotype including small and thinner cornea, abnormal iridocorneal angle, disorganized corneal stroma and neovascularization. In addition, the primary cilium ablation in NCC reduces the cell proliferation in a subpopulation of POM cells surrounding the RPE which are normally responsive to Ihh, the main Hh ligand expressed by endothelial cells of the choroid. Moreover, the primary cilium ablation in NCC decreases the expression of transcription factors associated with ASD in patients. These findings suggest that the primary cilium might contribute to the pathology of ASD, and associated complications like glaucoma.

## METHODS

### Mouse strains

Mouse strains *Ift88^tm1Bky^*, here referred to as *Ift88^fl/fl^*[84], *Wnt1-Cre* [45], *Gt(Rosa)26Sor^tm4(ACTB-tdTomato,-EGFP)Luo^/J* (*R26^mT/mG^*, Jackson Laboratories stock No 007676) [44], and *Gli1^tm2Alj^/J* (*Gli1-LacZ*, Jackson Laboratories stock No 008211) [56] were maintained on mixed C57Bl/6, FVB and 129 genetic backgrounds. *Gt(ROSA)26Sor^tm1(Sstr3/GFP)Bky^* mouse (here referred to as Sstr3::GFP mouse) [47] was maintained on a mixed CD1 background. *Ift88* conditional knockout (cKO) were generated by crossing *Wnt1-Cre*;*Ift88^fl/+^*males with *Ift88^fl/fl^* females. Since *Wnt1-Cre*;*Ift88^fl/fl^* mutants die at birth, all the study was conducted during the embryonic development. *Wnt1-Cre*;*Ift88^fl/+^*and *Ift88^fl/fl^* littermates were used as control. For time breeding, females were examined daily for vaginal plug. The day of the plug was considered embryonic day 0.5 (E0.5). All animal procedures were performed in accordance with the guidelines and approval of the Institutional Animal Care and Use Committee at Icahn School of Medicine at Mount Sinai, and at the University of Pennsylvania.

### Histology, electron microscopy and immunofluorescence

Mouse embryos at different gestational stages were removed from *Ift88^fl/fl^*females crossed with *Wnt1-Cre*;*Ift88^fl/+^*males and pictures were taken with a M80 stereomicroscope (Leica) equipped with a color IC80 HD camera (Leica). Half head or enucleated eyes from the embryos were fixed overnight in 4% paraformaldehyde (PFA) and embedded in paraffin for histological analysis. Hematoxylin and eosin (HE) staining was performed following standard procedures. Corneal stromal thickness was determined in the center of the cornea on HE sections from 3 mice per age.

Endomucin and TUJ1 staining were performed on fixed whole corneas as previously described [85, 86]. For immunofluorescence (IF) and X-gal staining on sections, half head and enucleated eyes from embryos were embedded and frozen in optimal cutting temperature compound (OCT Tissue-Tek, Sakura). For detection of the β-galactosidase activity by X-Gal staining, sections were fixed 10 min with 0.2% glutaraldehyde, 2 mM MgCl_2_ in PBS at room temperature, and then incubated overnight in a solution containing 1 mg/mL Xgal, 5 mM K_3_Fe(CN), 5 mM K_4_Fe(CN), 0.01% deoxycholate, 0.02% NP40 and 2 mM MgCl_2_ in PBS at 4°C. Sections were imaged with an AxioImager.Z2M microscope (Zeiss) equipped with an Axiocam 503 color camera (Zeiss). For immunofluorescent staining, sections were fixed 10 min with 4% PFA at room temperature. After blocking with 2% normal donkey serum (Jackson ImmunoResearch), 0.1% Triton in PBS for 30 min at room temperature, primary cilia were stained using a rabbit anti-Arl13b Ab (17711-1-AP, Protein Tech Group), pSmad2/3^+^ cells were stained using a goat anti-pSmad2/3 Ab (sc11769, Santa Cruz) and proliferative cells were stained using a rabbit anti-Ki67 Ab (ab15580, Abcam). Nuclei were counterstained with DAPI. Confocal pictures were acquired with a Leica DM6000.

For transmission electron microscopy (TEM), embryonic eyes were processed as previously described [87]. Briefly, samples were fixed with 1% PFA and 3% glutaraldehyde in 0.1 M sodium cacodylate buffer, post-fixed with 1% osmium tetroxide, embedded in Epon (Electron Microscopy Sciences) and stained with uranyl acetate and lead citrate after sectioning. Sections were imaged with Hitachi H7650 or S4300 microscopes.

### SAG stimulation

Enucleated eyes from P0 mice were cut in half on the sagittal axis and the lens and the retina were removed. Eyeball pieces were incubated for 5h in keratinocyte-SFM with or without 100 nM SMO agonist (SAG, MilliporeSigma) at 37°C in 5% CO2. Tissues were fixed 12 min with 4% PFA in PBS, incubated 12 min with 5% Triton in PBS and then primary cilia and SMO were stained using a mouse anti-acetylated tubulin Ab (6-11B-1, Sigma-Aldrich) and a rabbit anti-SMO Ab (1:100, ABS1001 MilliporeSigma), respectively. Whole-mount eyeball tissues were imaged using a Zeiss LSM800 confocal. Cilia and SMO staining were also performed on ocular tissues from E14.5 embryos, without prior incubation with or without SAG.

### *Ex vivo* imaging

Confocal imaging was performed on whole eyes from E18.5 *Wnt1-Cre*;*Ift88^fl/fl^*;*R26^mT/mG^*and *Wnt1-Cre*;*Ift88^fl/+^*;*R26^mT/mG^*embryos as previously described [87]. Live GFP and tdTomato fluorescences were recorded by a Zeiss LSM880 confocal microscope, using a 40X water immersion objective lens and used to scan a 0.05 mm^2^ field of view to study the keratocyte organization and 0.005 mm^2^ field of view to quantify the cytoplasmic processes. For the keratocyte organization, serial optical sections were acquired in 0.45 μm steps, through the entire cornea, from the corneal epithelium to the lens epithelium (49-89 μm per z-stack). For the quantification of cytoplasmic processes, serial optical sections were acquired in 0.2 μm steps, from the last epithelial cell layer before the corneal stroma to the first layer of keratocytes (∼5 μm per z-stack).

Quantification of the protrusions in Wnt1-Cre-positive keratocytes was done by counting cytoplasmic processes at the interface between corneal epithelium and stroma. From the acquired z-stacks, images from the first one where green fluorescent cytoplasmic processes are visible to the last one before keratinocyte cell bodies are imaged were selected and a maximum intensity projection was done with Fiji [53]. For each mouse, cytoplasmic processes were manually counted in 4 distinct areas in the center of the cornea and expressed as number of cytoplasmic processes per 100 μm^2^. Keratocyte organization in the stroma was assessed by quantifying the amount and the average size of extracellular spaces in the first 1/10 of the z-stack just under the corneal epithelium, corresponding to the anterior part of the stroma, and the last 1/10 of the z-stack just before the corneal endothelium, corresponding to the posterior part of the stroma (2.25-4.5 μm thickness/stack). After segmentation of the stacks with Ilastik [52], the amount and average size of extracellular spaces were quantified using Fiji (**Supplement Figure 2**).

For visualization of the blood vessels at the corneal periphery, the microscope field was centered on the major arterial circle which was considered as the border between the cornea and the sclera (**Supplement Figure 5**). Serial optical sections were acquired from the epithelium to the presumptive iris (49 to 109 μm per z-stack). After maximum intensity projections, the corneal area covered by blood vessels and the percentage of corneal and scleral thicknesses with blood vessels were quantified with Fiji.

### Intravital imaging

Intravital imaging of cilia in the mouse cornea was performed with an Olympus FV1200MPE microscope, equipped with a Chameleon Vision II Ti:Sapphire laser. Mice were anaesthetized with intraperitoneal injection of ketamine and xylazine (15 mg/mL and 1 mg/mL, respectively in PBS). A custom made stage was used to immobilize the mouse head and expose the eye globe. Mice were then placed under the microscope onto a heating pad and kept anesthetized with a continuous delivery of isoflurane through a nose cone (1% in air). A laser beam tuned at 930nm was focused through a 25X water immersion objective lens (XLPlan N, N.A.1.05; Olumpus USA) and used to scan a 0.5 mm^2^ field of view. Serial optical sections were acquired in 2-3 μm steps, starting from the surface of the eye and capturing the entire thickness of the cornea (epithelium ∼40 μm, stroma ∼80 μm).

### Cell sorting

#### Unstained cells

At E18.5, eyes from *Wnt1-Cre*;*Ift88^fl/+^*;*R26^mT/mG^*and *Wnt1-Cre*;*Ift88^fl/fl^*;*R26^mT/mG^* embryos were enucleated. After removing of the lens and retina from the eyeball, remaining tissues were digested by 8 mg/mL collagenase (Sigma) in keratinocyte-SFM (Thermo Fischer) for 1h at 37°C under stirring. Isolated cells were centrifuged for 5 min at 1800 rpm and re-suspended in sorting buffer (2% FBS in PBS) for cell sorting with a BD FACSAria (BD Biosciences).

#### Stained cells

Isolated cells from E18.5 *Wnt1-Cre*;*Ift88^fl/+^*;*R26^mT/mG^*and *Wnt1-Cre*;*Ift88^fl/fl^*;*R26^mT/mG^* embryonic scleras were incubated 45 min with an anti-CD31 Ab (BD Biosciences, 1:500), washed, and then incubated 30 min with an Alexa Fluor 647 donkey anti-rat secondary Ab. After washing, cells were re-suspended in sorting buffer for cell sorting with a BD FACSAria (BD Biosciences).

### Quantitative RT-PCR

Total RNA from the cornea dissected just under the iridocorneal angle, neural retina, the posterior half of the eyeball without the neural retina (corresponding to the sclera and the RPE layer) or sorted cells were extracted using RNeasy microkit (Qiagen) according manufacturer instructions. One hundred nanogram of RNA were reverse transcribed as previously described [87]. Real-time PCR was performed in triplicate using Maxima SYBR Green Master Mix (Thermo Fisher) on ABI PRISM 7900HT (Applied Biosystems). PCR program consisted of 40 cycles of 95°C for 15 s, 55°C for 15 s and 72°C for 30 s. Data were analyzed using the 2-ΔCt method, with *Gapdh* transcript as a reference. Primer sequences are listed in **Table 1**.

**Table 1.**
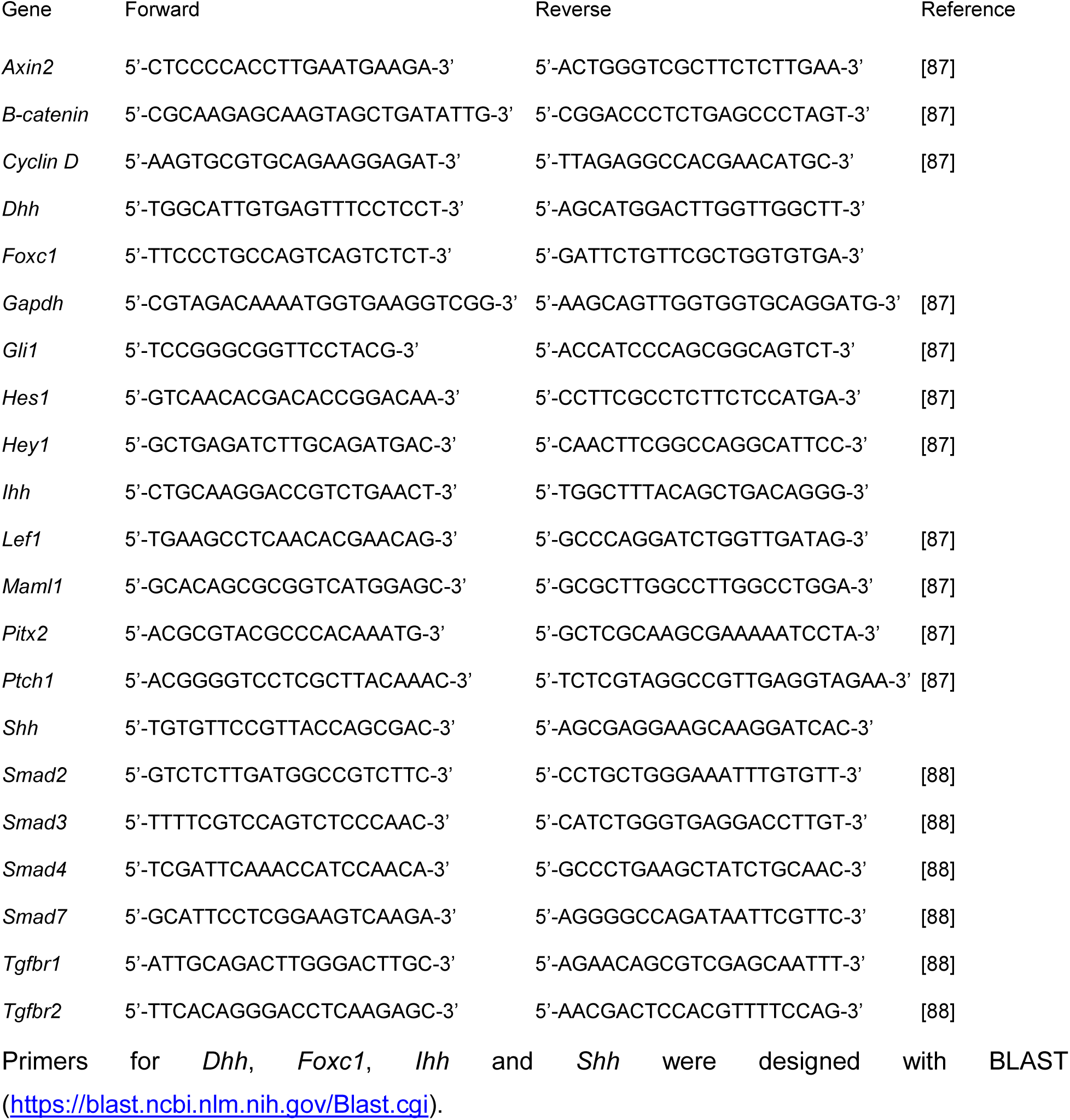
RT-qPCR primers

### Statistical analysis

Data are presented as mean ± SEM. Student’s *t*-tests were performed with Excel 2017 (Microsoft) and one-way ANOVA with post-hoc Tukey HSD test was performed with the online web statistical calculators https://astatsa.com/. A *P* value < 0.05 was considered significant.

## AUTHOR CONTRIBUTIONS

C.P. and C.I. conceived and designed the study. C.P., P.L., P.R. and C.I. performed the experiments, analyzed and interpreted the data. C.I., P.L. and P.R. provided funding for the project. C.P. and C.I wrote the manuscript.

## ACKNOWLEDGEMENTS

We thank N. Tzavaras (Microscopy CoRE, Icahn School of Medicine at Mount Sinai), V. Nair and N. Marjanovic (qPCR CoRE, Icahn School of Medicine at Mount Sinai), and C. Bare and P.-Y. Kuo (Flow cytometry CoRE, Icahn School of Medicine at Mount Sinai) for their technical support. We thank all Mlodzik and Iomini lab members and the Wilmer Cornea Group members for helpful inputs and discussion, and Rebecca Sherburn, and James Foster for critical comments on the manuscript. This work was supported by National Institutes of Health grant EY022639 to C.I. and EY030599 to P.R., and the Research to Prevent Blindness Dolly Green Special Scholar Award to C.I.

