## Supplemental data for "Primary cilia deficiency in neural crest cells causes Anterior Segment Dysgenesis"

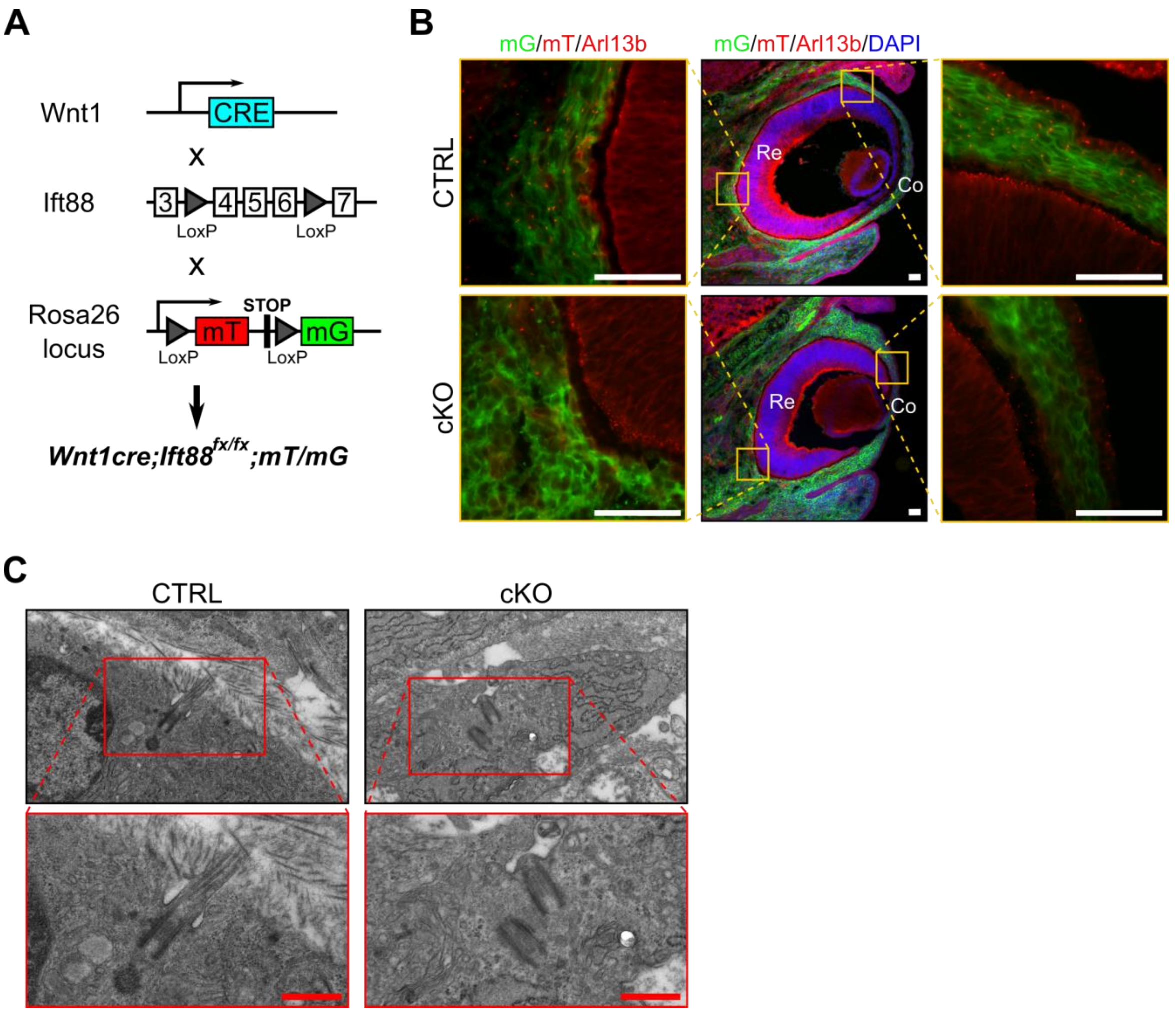


**Supplement Figure 1: Genetic deletion of Ift88 in NCC leads to primary cilium ablation in NCC.** (**A**) Breeding strategy to generate NCC ciliary mutant and visualize Cre expression. (**B**) Representative eye sections of control and cKO embryos at E14.5. NCC express the mG reporter (green cells) whereas cells from other embryonic origin express the mT reporter (red cells). Primary cilia were stained with an anti-Arl13b Ab and appear as small red rods. Scale bar, 50 µm; Co, cornea; Re, retina. (**C**) Representative pictures of primary cilia in the corneal stroma at E17.5. In contrast to control, primary cilia do not assemble in cKO embryos. Scale bar, 0.5 µm.

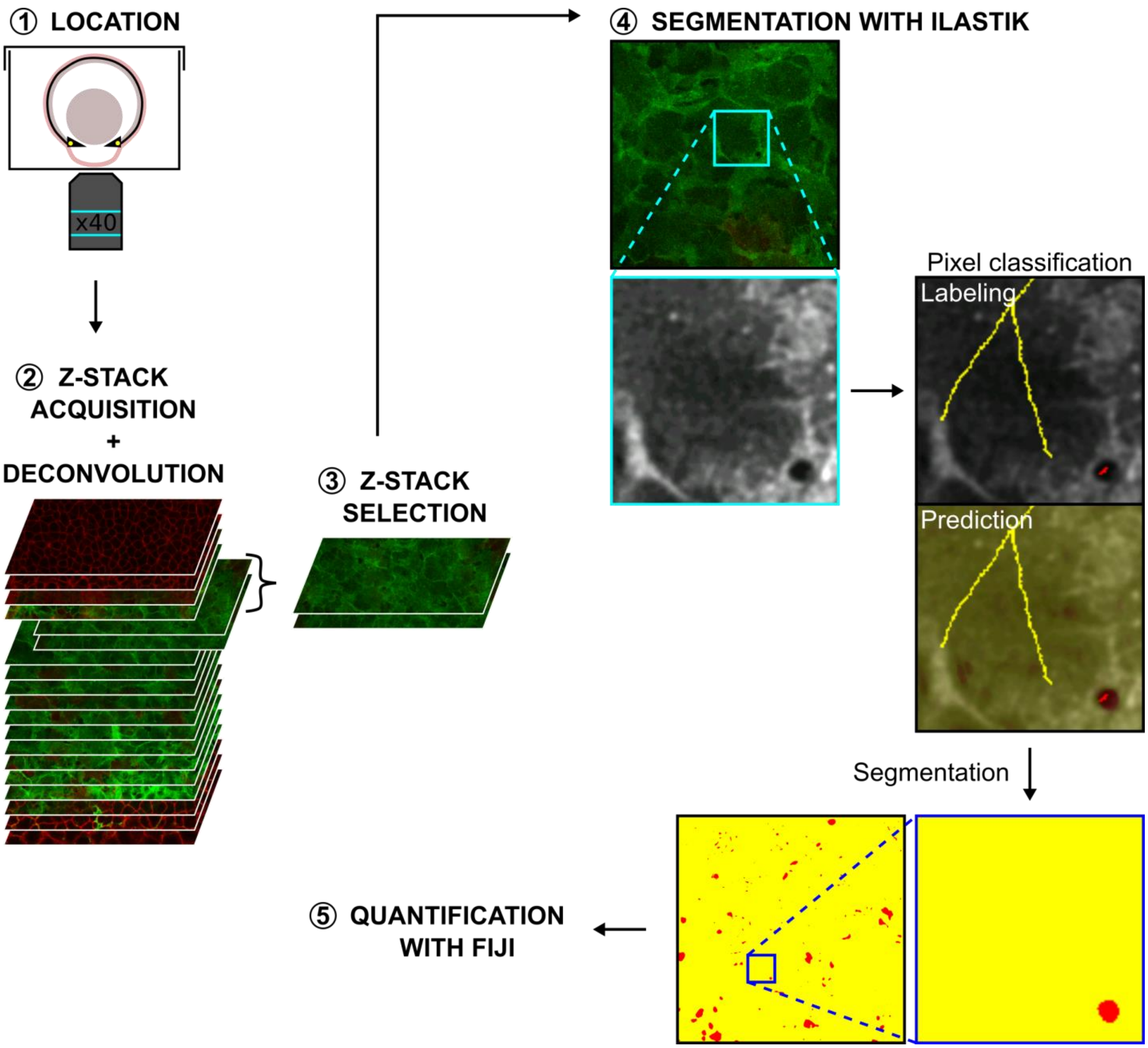


**Supplement Figure 2: Segmentation of the extracellular spaces with Ilastik to study corneal stroma organization. 1.** Enucleated eye was placed facing down in a glass-bottom microwell dish. **2.** A z-stack was acquired from the corneal epithelium to the lens epithelium by confocal microscopy, and then deconvoluted with AutoQuant X3. **3.** z-sections corresponding to the area of interest were selected. **4.** After uploading the z-sections in Ilastik, a pixel classification was done to train the classifier to recognize pixels belonging to the cells or to the extracellular spaces. For this, a trained user labeled pixels with a different color for each class (here, cells in yellow and extracellular spaces in red). Boxed region indicates the area shown at higher magnification below. Quality of the segmentation was checked with the prediction. These steps were repeated on different z-sections and as many times as necessary to obtain a good segmentation. Finally, the segmentation is completed. **5.** Segmented masks were then exported and the quantification of the amount and the average size of extracellular spaces were performed with Fiji.

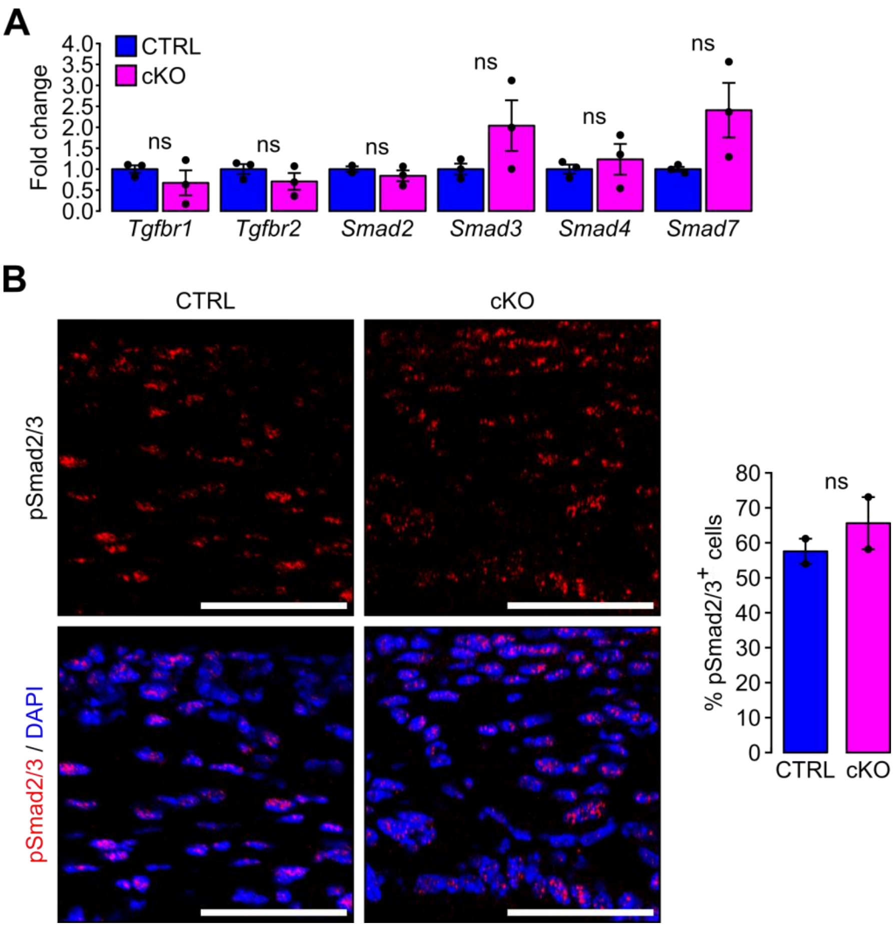


**Supplement Figure 3: TGFβ signaling pathway is not affected by the primary cilium ablation in NCC.** (**A**) Gene expression was evaluated in corneas of E18.5 embryos by RT-qPCR. Data are mean ± SEM. Statistical significance was assessed using two-tailed Student's *t*-test (n=3 embryos/group). ns, non-significant, P ≥ 0.05. (**B**) Immunostaining of phosphorylated Smad2/3 (pSmad2/3) in the corneas of E18.5 embryos. The proportion of pSmad2/3$^+$ cells in the cornea is not affected by the primary cilium ablation in NCC. Scale bar, 50 µm. Data are mean ± SEM. Statistical significance was assessed using two-tailed Student's *t*-test (n=2 embryos/group). ns, non-significant, P ≥ 0.05.

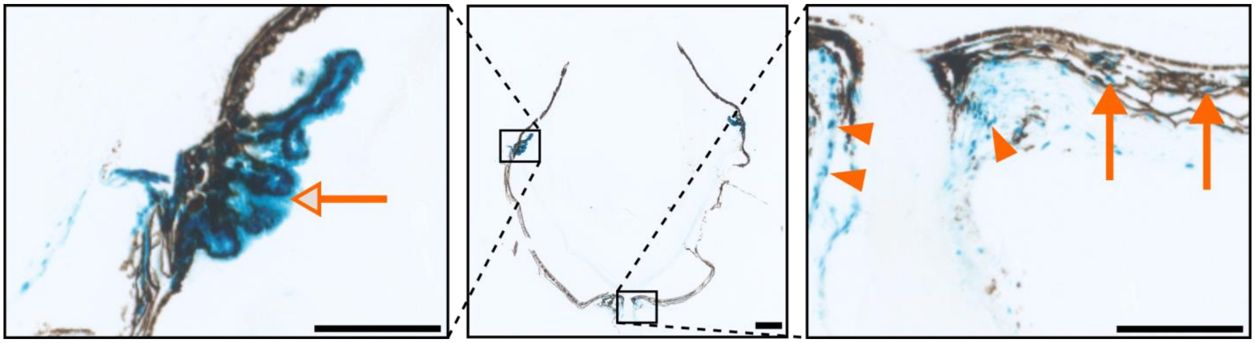

**Supplement Figure 4: Hh activity in the eye at adulthood.** Hh activity assessed by Gli1-LacZ staining at adulthood (5 months). Hh signaling remains active in the ciliary body (empty arrow), in the choroid (orange arrows) and around the optic nerve (arrowheads). Scale bar, 250 µm.

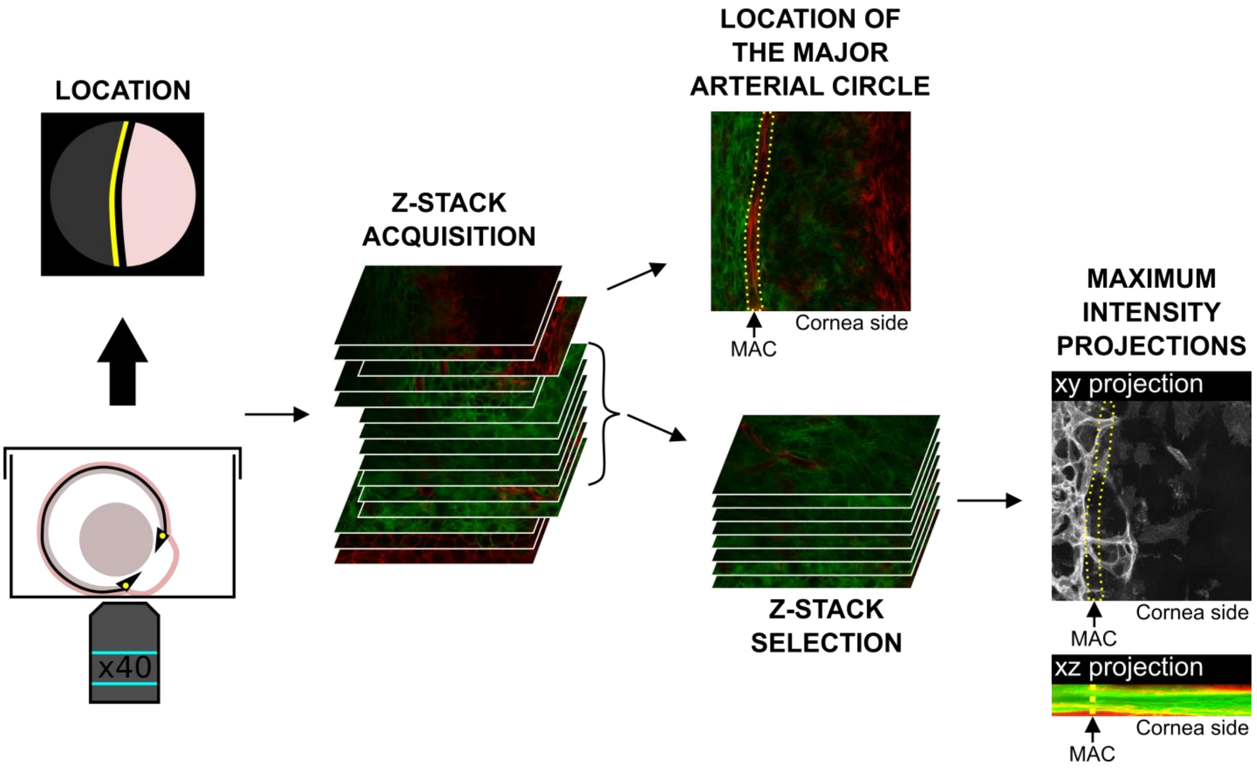


**Supplement Figure 5: Live imaging of the cornea boundary area.** Enucleated eye was placed facing down in a glass-bottom microwell dish. The observed field was centered on the major arterial circle (yellow line). A z-stack was acquired from the corneal epithelium to the presumptive iris by confocal microscopy (49 to 109 µm per stack). and then deconvoluted with AutoQuant X3. z-sections corresponding to the area of interest were selected. The major arterial circle (yellow dotted line) is the reference to define the boundary between the cornea and the sclera. Quantifications of the corneal area covered by blood vessels and the percentage of corneal and stromal thickness with blood vessels were done with Fiji on the maximum intensity projections. MAC, major arterial circle.

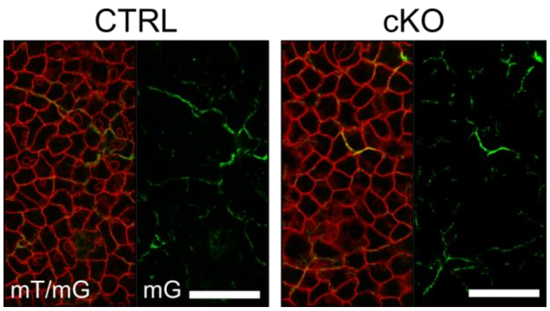


**Supplement Figure 6: Innervation of the corneal epithelium.** Representative confocal pictures of the epithelium in the center of the cornea of E18.5 embryos imaged on live tissues using the mT/mG reporter. Corneal epithelial cells are visualized with the tdTomato fluorescent reporter (mT) and the sensory nerves with the GFP fluorescent reporter (mG). Corneal epithelia of both genotypes are innervated. Scale bar, 50 µm.
